# Balancing p38 MAPK Signaling with Proteostasis Mechanisms Supports Tissue Integrity during Aging in *C. elegans*

**DOI:** 10.1101/2022.07.12.499791

**Authors:** Wang Yuan, Yi M. Weaver, Svetlana Earnest, Clinton A. Taylor, Melanie H. Cobb, Benjamin P. Weaver

## Abstract

Like other p38 MAPKs, *C. elegans* PMK-1 is activated by phosphorylation during stress responses and inactivated by phosphatases. PMK-1 initiates immune response and blocks development when hyperactivated. Here we show that PMK-1 signaling is essential for tissue homeostasis during aging. Loss of PMK-1 accelerates progressive declines in neuronal integrity and lysosome function compromising longevity. Enhancing p38 signaling with caspase cleavage-resistant PMK-1 protects lysosomal and neuronal integrity extending a youthful phase. The cleavage-resistant PMK-1 mutant behaves oppositely to the *pmk-1* null in regulating both transcriptional and protein degradation programs supporting tissue homeostasis. PMK-1 activates a complex transcriptional program and requires UNC-62 (MEIS), FOS/JUN, and DAF-16 (FOXO) transcription factors to regulate lysosome formation which is both cell autonomous and non-cell autonomous. We show that during early aging the absolute phospho-p38 amount is nearly constant but maintained at a small percentage of total p38. The reservoir of non-phospho-p38 diminishes during early aging to enhance signaling without hyperactivation. CED-3 caspase cleavage limits phosphorylated PMK-1 and truncated PMK-1 is rapidly degraded by the proteasome. Modulating phospho-p38 ratio confers dynamic control for tissue-homeostasis without activating stress response to support longevity.

## Introduction

The p38 family consists of stress-sensing MAPKs that respond to diverse stimuli including chemical and physical stressors, microbial infections, as well as cytokine cues (Canovas and Nebreda 2021). Licensing of p38 MAPK signaling pathways following physiological stressors relies on dual phosphorylation of the highly conserved TGY motif in the activation loop by upstream activating kinases. Recently, it was shown that both pathogenic and non-pathogenic stresses initiate multimerization of the membranous Toll/interleukin-1 receptor domain protein (TIR-1) receptor ultimately leading to activation of the PMK-1 immunity program in intestinal cells (Peterson et al. 2022). Balancing this activation, phosphatases including dual specificity phosphatases (DUSPs) dephosphorylate the TGY motif, making for cycles of activation and deactivation (Raman et al. 2007). In the case of *C. elegans* PMK-1 p38 MAPK, VHP-1 is the specific DUSP that limits PMK-1 phosphorylation (Kim et al. 2004; Mizuno et al. 2004).

Stress response pathways have complex effects on aging. In *C. elegans,* the p38 MAPK branch supports innate immunity in response to stress throughout life. Activation of SKN-1 (NRF) by PMK-1 is required to ensure longevity by maintaining proper redox balance (Inoue et al. 2005; Tullet et al. 2008). Loss of the PMK-1-dependent pathogen-response program results in immunosenescence with aging, thereby shortening life span when animals are exposed to pathogenic bacteria (Troemel et al. 2006; Youngman et al. 2011).

Caspases have been identified as key regulators of non-apoptotic cell fate during metazoan development. Muscle cell differentiation in mice requires chromatin reorganization by elimination of PAX7 and SATB2 by caspases-3/7 (Dick et al. 2015; Bell et al. 2022). Pluripotency potential is limited in an epidermal stem cell lineage in *C. elegans* by CED-3 caspase and UBR-1 E3 ligase-mediated elimination of LIN-28 (Weaver et al. 2014; Weaver et al. 2017). Asymmetric cellular fates for neuroblasts in *C. elegans* is determined by CED-1 (MEGF10 receptor) and CED-3 caspase antagonizing PIG-1 (MELK)-dependent mitotic potential (Mishra et al. 2018). Intestinal progenitor cell quiescence in flies is determined by Dronc caspase regulation of Notch signaling (Arthurton et al. 2020). These findings underscore the contribution of caspases working outside of cell death to ensure cell fate decisions.

We recently reported that CED-3 caspase antagonizes a PMK-1-dependent anti-microbial function to promote post-embryonic development in *C. elegans* (Weaver et al. 2020). PMK-1 hyperactivation is critical for survival in the presence of pathogens or abiotic stressors but can be detrimental leading to stalled development (Mizuno et al. 2004; Kim et al. 2016; Foster et al. 2020; Weaver et al. 2020). The VHP-1 phosphatase is an essential gene underscoring the importance of keeping stress-responsive phosphorylated PMK-1 (pPMK-1) in check (Mizuno et al. 2004). Altogether, these findings suggest that p38 MAPKs may have broader functions in integrating cross-talk between stress responses and developmental programs.

Proteostasis involves a complex network of factors ensuring native protein functions while also mitigating aberrant protein functions and this network is progressively challenged during aging (Ben-Zvi et al. 2009; Walther et al. 2015; Visscher et al. 2016). Long-lived mutants have enhanced proteostasis control including clearance of misfolded proteins (Hipp et al. 2019). Proteostasis clearance pathways include the ubiquitin-proteasome system as well as the autophagy-lysosome pathway and disruption of these degradation systems compromises health and lifespan (Morimoto 2020). The proteostasis network is best understood for its role in mitigating stressful or pathological states such as the heat shock response which requires integration of signaling between cells to sense and respond accordingly (Morimoto 2020). Stress signaling to restore proteostasis is critical for health and longevity. However, the impact of proteostasis on cell signaling pathways and its effects on normal tissue aging remains a major area for further understanding.

## Results

### PMK-1 regulates lifespan and is modulated by CED-3 caspase cleavage

We previously identified *C. elegans* PMK-1 as a substrate of CED-3 caspase to limit stress responsiveness during larval development (Weaver et al. 2020). The CED-3 cleavage site is found among multiple p38 members from diverse phyla and located just beyond the core kinase domain at the beginning of the C-terminal extension (CTE), which is highly conserved throughout metazoans (Fig. 1A-B) and solvent accessible (Supplemental Fig. S1A-B). Additional putative caspase cleavage sites outside of the CTE for human p38γ are also solvent accessible (Asp46 and Asp101, Fig. 1B; Supplemental Fig. 1B). The CTE is important for promoting active MAPK conformations, suggesting that cleavage within this region will impair p38 function.

**Figure 1.**
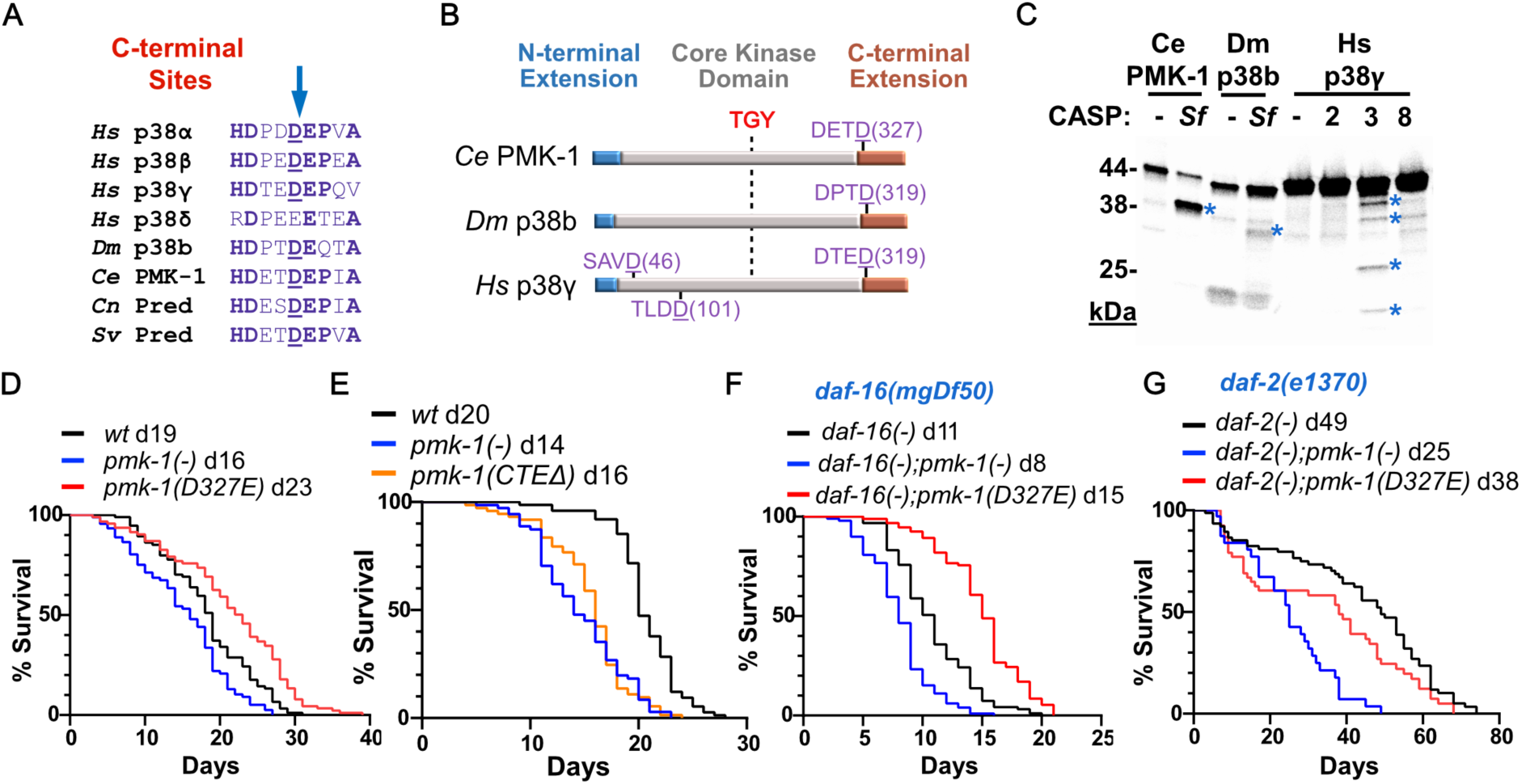
The p38 MAPK C-terminal region is conserved and caspase cleavage of PMK-1 controls longevity in *C. elegans.* (*A*) Local alignment of p38 homologs putative cleavage site (arrow) at C-terminal Extension. Predicted PMK-1 homologs (Pred). *Strongylus vulgaris, Sv; Caenorhabditis nigoni, Cn; Caenorhabditis elegans, Ce; Drosophila melanogaster, Dm; Homo sapiens, Hs.* (*B*) Relative positions of p38 dual phospho-activation sequence (TGY) and predicted cleavage sites (purple) (*C*) *In vitro* caspase cleavage assay. ^35^S-labelled proteins incubated without (-) or with indicated active caspase. *Sf,* Sf21 insect cell extracts with active caspase; Human caspases-2, 3, and 8. Asterisks indicate cleavage products. (*D-G*) Aging assays of outcrossed strains on normal OP50/NGM media without FUDR. Median survival days indicated for each strain next to genotypes. Replicates and statistics (Supplemental Fig. 1).

Using *in vitro* cleavage assays, we found that in addition to *C. elegans* PMK-1, *Drosophila* p38b and human p38γ are candidate caspase substrates (Fig. 1C). In this assay, *C. elegans* PMK-1 was cleaved by insect caspase in a manner identical to cleavage by CED-3. We studied *C. elegans* PMK-1 further to understand how p38 signaling broadly impacts physiology in normal feeding conditions.

In previous studies, loss of *pmk-1* function has been shown to compromise longevity during exposure to pathogenic bacteria but no effect on aging was revealed on non-pathogenic food when germline proliferation was blocked by treatment with FUDR (Troemel et al. 2006; Wu et al. 2019). We tested the effects of *pmk-1* null mutants on aging when fed normal food (no FUDR) allowing for normal germline proliferation. We found that loss of *pmk-1* function had shortened adult lifespan under normal dietary conditions (Fig. 1D). Our results differ significantly from the previous findings when FUDR was used to stop germline proliferation. This difference is consistent with the tradeoff of fecundity versus proteostasis in determining lifespan (Labbadia and Morimoto 2015; Hipp et al. 2019).

To abolish caspase cleavage of PMK-1, we generated a *pmk-1(D327E)* cleavage-resistant mutation in the endogenous locus using CRISPR mutagenesis and outcrossed multiple times (Materials and methods; Supplemental Table S1-S3). Changing aspartate to glutamate in this position abolishes cleavage but preserves charge and is naturally occurring in some p38 and many MAPK homologs (Fig. 1A). In contrast to our findings of short-lived *pmk-1* null mutants (Fig. 1D), the caspase cleavage-resistant *pmk-1(D327E)* mutants had an extension of lifespan (Fig. 1D; Supplemental Fig. S1C; Supplemental Table S4). Loss of the PMK-1 C-terminal extension (CTEΔ) causes considerably shortened lifespan similar to the *pmk-1(-)* suggesting that the cleaved protein is not functional (Fig. 1E). In addition to acting on PMK-1, CED-3 caspase has many other critical functions including germline apoptosis during adulthood. Thus, it is not surprising that we observed *ced-3* null mutants had diminished life span (Supplemental Fig. S1C).

Insulin is critical for metabolic adaptation and basal stress resistance, functioning as a major determinant of longevity in nematodes (Lin et al. 1997; Ogg et al. 1997), flies (Clancy et al. 2001; Tatar et al. 2001) and mice (Bluher et al. 2003; Holzenberger et al. 2003). To determine the relationship between PMK-1 signaling and insulin-dependent longevity, we tested how *pmk-1* null and *pmk-1(D327E)* mutations impact the lifespans of *daf-16* (FOXO transcription factor) and *daf-2* (insulin receptor) mutants. We found that the cleavage-resistant *pmk-1(D327E)* mutation restored a more normal lifespan in *daf-16-*deficient mutants, whereas *pmk-1* null mutants further reduced lifespan in animals with compromised *daf-16* function (Fig. 1F; Supplemental Fig. S1D; Supplemental Table S4). Extending lifespan of *daf-16* mutants by *pmk-1(D327E)* mutation suggests that a key sub-set of *pmk-1-*dependent genes act through additional transcriptional pathways supporting longevity.

Surprisingly, both *pmk-1* null mutation and the cleavage-resistant mutation reduced longevity of *daf-2* deficient mutants (Fig. 1G). The reduction of *daf-2* longevity by *pmk-1* null mutation is consistent with previous work demonstrating that *daf-2-*enhanced pathogen resistance required intact *pmk-1* (Troemel et al. 2006). We find that the *pmk-1(D327E)* cleavage-resistant mutation compromised *daf-2* longevity suggesting that either the heightened PMK-1 function can partially restore *daf-2* signaling or promote a competing gene regulatory program.

### Phospho-p38 pool is strictly maintained in animals and non-phospho-p38 pool is dynamic throughout aging

We then considered how CED-3 cleavage impacts PMK-1 protein expression. To monitor protein expression during development and aging, we used CRISPR mutagenesis to add an HA tag at the N-terminus of endogenous PMK-1 (wt, Fig. 2A). We also added the cleavage-resistant D327E mutation to the endogenous locus to compromise CED-3 cleavage *in vivo* (DE, Fig. 2A). Adding tags at the N-terminus of PMK-1 does not affect function as determined by induction of a PMK-1-dependent reporter and comparable expression profiles (Supplemental Fig. S2A).

**Figure 2.**
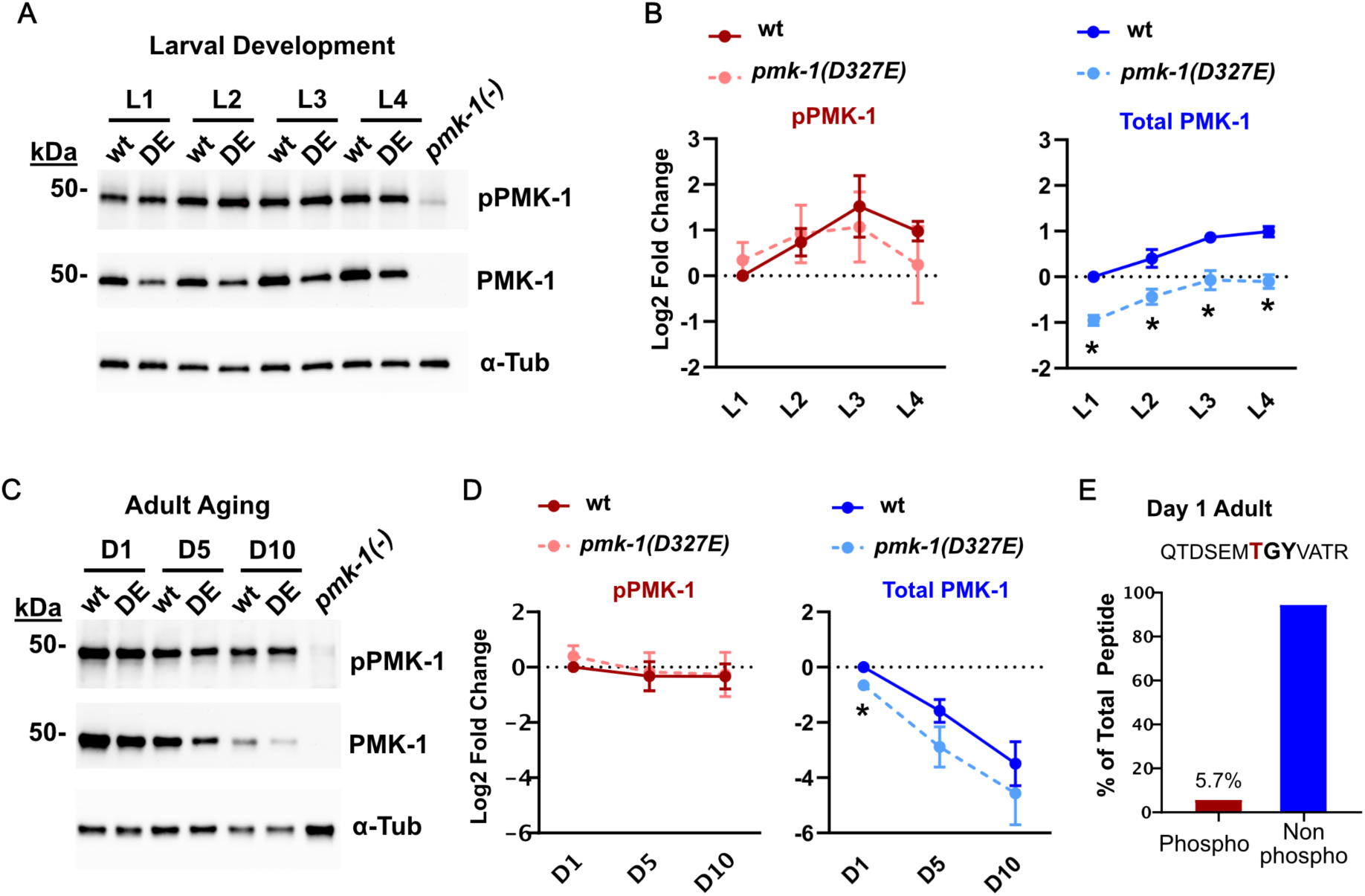
Ratio of phospho-PMK-1 to non-phospho-PMK-1 changes during development and aging. (*A-B*) Expression and quantitation of pPMK-1 and total PMK-1 protein through post-embryonic development. Endogenous *pmk-1* locus tagged with HA with either Asp327 (wild-type) or cleavage-resistant mutation Glu327 (DE) using CRISPR mutagenesis. Mean values of 3 independent biological replicates were plotted as Log2 fold change normalized to α-tubulin and relative to wt L1 stage (set to 1.0). (*C-D*) Expression and quantitation of pPMK-1 and total PMK-1 protein throughout aging. Mean values of 3 independent biological replicates were plotted as Log2 fold change normalized to α-tubulin and relative to wt Day 1 adults (set to 1.0). *(E)* Mass spectrometric determination of phospho-peptide to non-phospho-peptide for the dual phosphorylation site (TGY)-containing peptide.

Interestingly, in wild-type animals, we found that the amounts of pPMK-1 peaked in mid-larval development and were relatively constant throughout adulthood (pPMK-1, Fig. 2B,D) whereas total PMK-1 protein modestly increased by the end of larval development (Total PMK-1, Fig. 2B) but markedly diminished with aging (Total PMK-1, Fig. 2D). It is intriguing that the pool of phosphorylated PMK-1 is maintained fairly constant during aging but non-phosphorylated PMK-1 levels are dramatically downregulated during aging at a time when PMK-1 function is critical to support longevity. Using mass-spectrometry, we find that the phosphorylated TGY-containing peptide for PMK-1 (QTDSEMTGYVATR) is limited to about 1 out of 20 peptides for that sequence indicating that phospho-PMK-1 levels are kept limited at early adulthood (Fig. 2E).

Surprisingly, the amount of phosphorylated PMK-1(D327E) was similar to phosphorylated wild-type PMK-1 (pPMK-1), but total PMK-1(D327E) protein was reduced by ∼50% at each stage of larval development (Fig. 2A-B; Supplemental Fig. S2B) and throughout adulthood (Fig. 2C-D; Supplemental Fig. S2C). Steady state *pmk-1* mRNA was comparable between wild-type and the cleavage-resistant mutant (Supplemental Fig. 2D). Moreover, the decrease in PMK-1 protein resulting from D327E mutation was contrary to our prediction that failure to cleave PMK-1 would result in PMK-1 accumulation. Therefore we further tested the possibility that down-regulating the total PMK-1 protein was an adaptive mechanism to the keep the amount of phospho-PMK-1 constant in the absence of caspase cleavage.

### Phospho-p38 pool is limited by caspase with truncated PMK-1 degraded by proteasome

Finding that animals tightly regulate phospho-PMK-1 levels throughout adulthood suggests robust regulatory mechanisms of PMK-1 *in vivo* during aging. To overcome this, we generated an auxin-controlled overexpression system for transient overexpression. We used CRISPR mutagenesis to add a second copy of PMK-1 in a MosI site (materials and methods). The PMK-1 was tagged with HA and driven by a constitutive promoter (*eft-3*). For wild-type overexpression (WT O/E), we used full length PMK-1 open reading frame with intact exons and introns. For cleavage-resistant PMK-1 (DE O/E), we took the same ORF and made a D327E mutation. To overcome adaptive compensation, we maintained these strains on auxin plates to prevent overexpression. This tool allowed us to test whether the decreased PMK-1(D327E) total protein (Figs. 2B, 2D) was an adaptive mechanism for enhanced PMK-1 signaling. Placing synchronous L1 animals on normal food without auxin to allow overexpression, we found that the cleavage-resistant mutation was able to increase the amount of phosphorylated PMK-1 about 10-fold compared to the wild-type overexpression (Fig. 3A-B; Supplemental Fig. S3A). This result confirmed that animals tightly regulate the amount of phosphorylated PMK-1 and down-regulating total PMK-1 protein is another mechanism in addition to dephosphorylation to compensate for excessive phosphorylated PMK-1.

**Figure 3.**
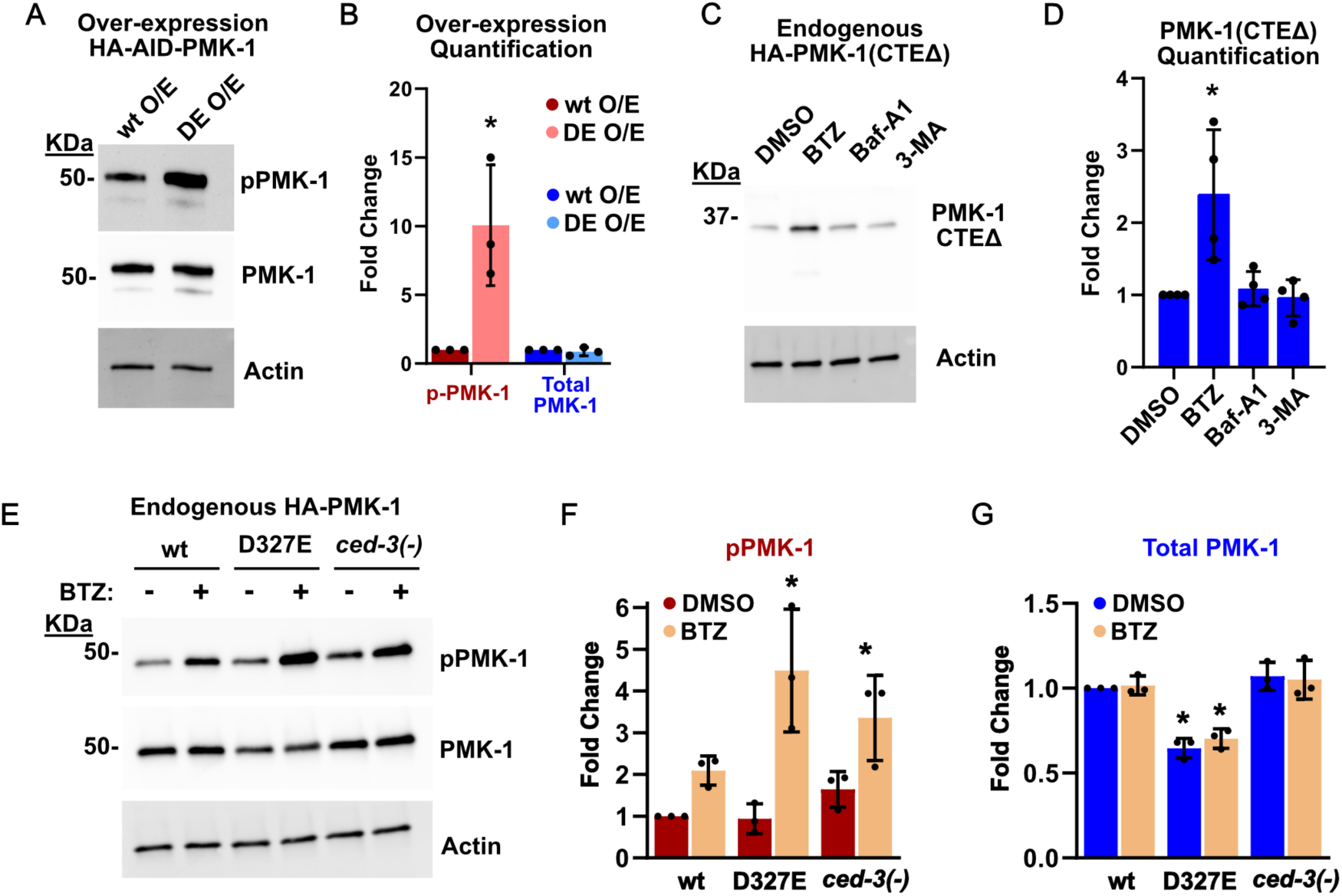
CED-3 caspase and proteasome limit phospho-PMK-1 levels. (*A-B*) Western Blot analysis of pPMK-1 and total PMK-1 from overexpressed wild-type PMK-1 (wt) or overexpressed cleavage-resistant PMK-1 (DE). Bar graph shows mean fold change of 3 biological replicates following treatments normalized by actin and relative to wt overexpression. (*C*) Effects of protein degradation pathways on expression of truncated PMK-1. CRISPR mutagenesis was used to remove the C-terminal domain from the endogenous *pmk-1* locus. Western blot analyses of truncated PMK-1 levels following treatment with control (DMSO); proteasome inhibitor, Bortezomib (BTZ); lysosome acidification inhibitor, Bafilomycin A1 (Baf-A1); and autophagy inhibitor, 3-methyladenine (3-MA). (*D*) Quantitation of three independent experiments. Protein levels following treatments normalized by actin and relative to DMSO treatment. (*E-G*) Western blot analysis and quantitation of endogenous pPMK-1 and total PMK-1 from wild type, cleavage-resistant PMK-1 (D327E) and caspase *ced-3(-)* mutant following Bortezomib treatment.

We did not observe a cleavage product *in vivo* suggesting possibly rapid degradation. To understand the effect of removing the C-terminus, we used CRISPR mutagenesis to generate a truncated HA-tagged PMK-1 at Asp327 in the endogenous locus (CTEΔ). We tested two lines of the CTEΔ mutation and observed only trace levels of the expected ∼38 kDa truncated protein (Supplemental Fig. S3C). The lack of CTE expression was not due to reduced mRNA (Supplemental Fig. S3B). To rule out solubility issues, we expressed wild-type PMK-1, PMK-1(D327E) and PMK-1(CTEΔ) proteins *in vitro* using reticulocyte lysates and found all three versions of PMK-1 were synthesized to comparable extents in this heterologous system (Supplemental Fig. S3C). Incubating these proteins for 48 hours at 20°C and 37°C did not result in loss of protein or accumulation of insoluble aggregates (Supplemental Fig. S3C) suggesting no gross differences in solubility and the truncated protein is likely targeted for degradation *in vivo*.

To identify the clearance pathway that degrades truncated PMK-1, we tested inhibitors of proteasome (Bortezomib, BTZ), lysosome (Bafilomycin A1, Baf-A1), and autophagy (3-methyladenine, 3-MA) and found that blocking the proteasome significantly increased accumulation of truncated PMK-1 (CTEΔ, Fig. 3C-D). These findings indicated that the cleaved PMK-1 intermediates are rapidly degraded by the proteasome clearance pathway.

We then tested the effect of proteasome inhibition on accumulation of phosphorylated PMK-1 (pPMK-1). We found that inhibiting the proteasome in wild-type animals caused a two-fold increase in accumulation of pPMK-1 (Fig. 3E-F). Furthermore, animals with either the PMK-1(D327E) cleavage-resistant mutation or the *ced-3* caspase null mutation accumulated pPMK-1 by nearly two-fold more than wild-type animals (Fig. 3E-F; Supplemental Fig. S3D). Again, consistent with our finding that phospho-PMK-1 is maintained at a small fraction of total PMK-1 (Fig. 2E), we did not observe any significant increases in total PMK-1 (Fig. 3G). Because *pmk-1(D327E)* mutants have intact CED-3 caspase function and *ced-3* null mutants have wild-type PMK-1, our results suggest that CED-3 caspase targets phospho-PMK-1 to proteasome for degradation.

### PMK-1 regulates gene expression and degradation programs to support tissue homeostasis

To understand how *pmk-1* null mutation and the cleavage-resistant *pmk-1(D327E)* mutation inversely impact longevity, we examined their effects on global gene expression programs. Using mRNA-Seq, we examined global steady-state mRNAs of *pmk-1* null mutant compared to wild-type to identify *pmk-1-*dependent genes under normal feeding conditions (Fig. 4A; Supplemental Table S5).

**Figure 4.**
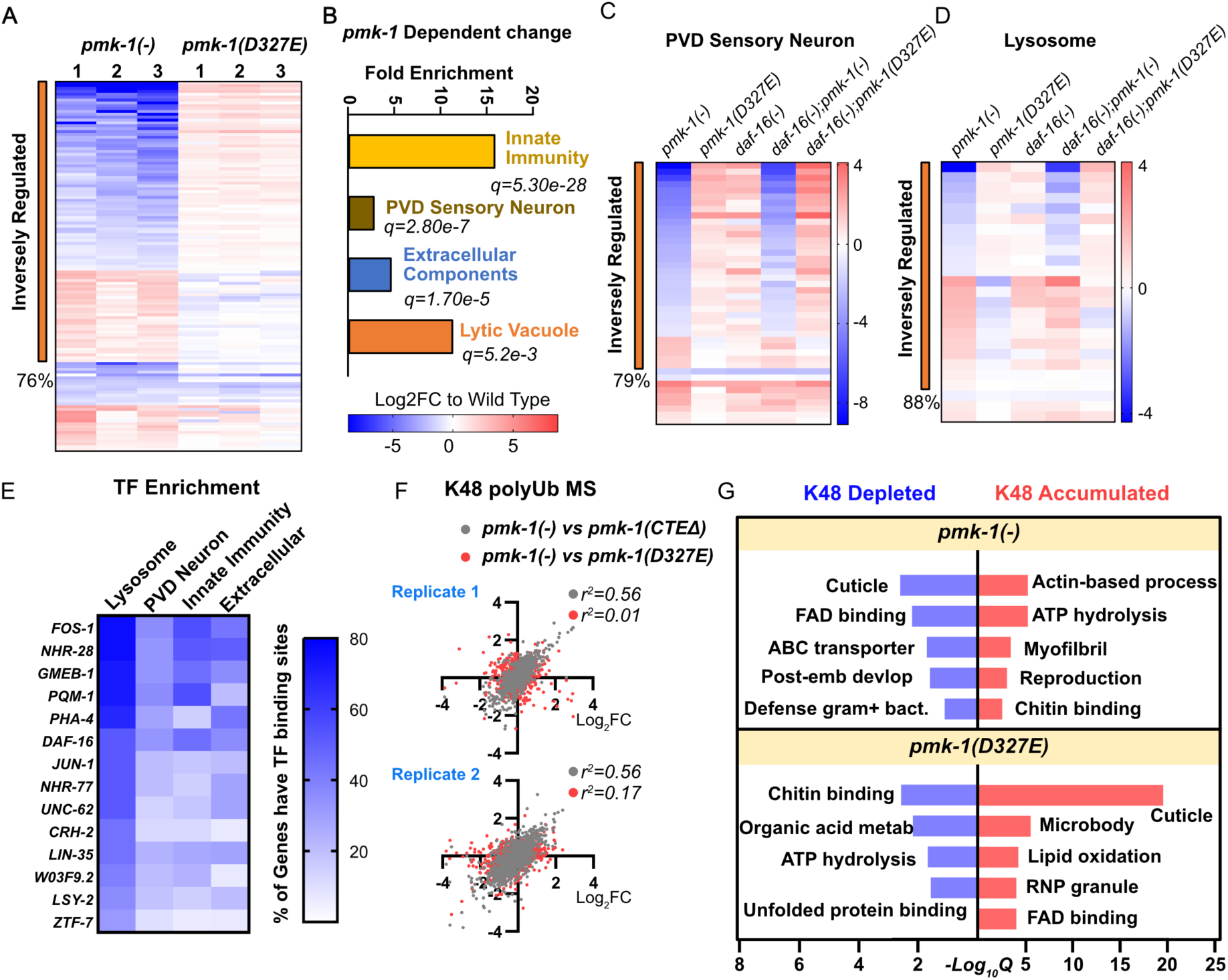
PMK-1 signaling regulated by caspase cleavage impacts gene expression and degradation programs regulating tissue maintenance. (*A*) mRNA-Seq and enrichment analysis to identify *pmk-1*-dependent genes in day 1 young adults. Both *pmk-1* null (-) and *pmk-1(D327E)* were compared to wild-type. Three biological replicates were analyzed and genes with false discovery rate < 0.1 for *pmk-1(-)* shown in heat map with Log2-fold change for each replicate. (*B*). Fold enrichment and -log10Q values of PMK-1 regulated genes (Gene Enrichment, Wormbase). (*C-D*) Heatmaps of *pmk-1-*dependent sensory neuron and lysosome genes for *daf-16* dependence. PMK-1-dependent genes clustered by inverse regulation and plotted with Log2-fold change. All mutants including *daf-16(mgDf50)* single and double mutants were compared to wild-type. (*E*) Frequency of transcription factor binding sites in proximal promoters of lysosomal, neuronal, innate immunity, and extracellular component genes. (*F-G*) Mass-spectrometric analyses of Lys48 (K48)-specific Ub linkages shows altered enrichments. Pearson’s test was used to determine correlation and *r^2^*values of variance. Wormbase enrichment analysis toolkit determined significant GO terms.

We also examined global steady-state mRNAs of *pmk-1(D327E)* mutant. In fact, we did observe that 76% of genes altered by the *pmk-1* null mutation were inversely regulated by the cleavage-resistant PMK-1(D237E) mutation (Fig. 4A).

We then performed enrichment analysis for genes altered by loss of *pmk-1*. Consistent with previous findings (Troemel et al. 2006; Youngman et al. 2011), loss of *pmk-1* resulted in a significant decline in innate immunity genes (Fig. 4B). Intriguingly, we also found altered expression of PVD-type sensory neuron genes, extracellular components, such as collagens and proteases, as well as lytic vacuole genes (Fig.4B). The expression of majority of these genes were reversed in mutants with the cleavage-resistant PMK-1.

We examined the PVD sensory neuron and lysosomal gene sets further for their effects on expression with and without functional *daf-16(FOXO).* Most genes altered by loss of *pmk-1* are also affected by loss of *daf-16,* suggesting PMK-1 signaling and DAF-16 co-regulate these genes. However, the inverse effects of *pmk-1(-)* versus *pmk-1(D327E)* for both PVD (Fig. 4C) and lysosomal genes (Fig. 4D) was not grossly affected by *daf-16* status, indicating there are other transcription factors in this regulatory network. Another *daf-16* mutation had comparable findings (Supplemental Fig. S4A-B). Importantly, loss of *pmk-1* resulted in dysregulation for both neuronal and lysosomal genes including some factors upregulated while others were down-regulated (Fig.4C-4D). We were particularly interested in these results given the importance that maintaining proper stoichiometry of organelle components has on maintaining proteostasis.

Based on our findings that *daf-16* was not strictly required for PMK-1 regulation of lysosomal and neuronal genes, we examined the proximal promoters of the inversely regulated genes to identify potential other transcription factor binding site enrichments (Fig. 4E; Supplemental Table S6). These enrichments were then compared to PMK-1-dependent PVD neuronal, innate immunity, and extracellular matrix genes (Fig. 4E). Altogether, our findings predict a model whereby PMK-1 p38 MAPK signaling can work with DAF-16 and a variety of additional transcription factors to regulate gene expression programs.

In addition to gene expression, we investigated how *pmk-1* null and *pmk-1(D327E)* impact protein clearance by the proteasome degradation pathway. We enriched for K48-linked poly-ubiquitin conjugated proteins following a brief treatment with Bortezomib to accumulate proteasome substrates. We found that total K48-linked protein accumulation was comparable amongst all strains (Supplemental Fig. S4C-E). We then performed K48-specific immunoprecipitation and identified proteins pulled down by mass-spectrometry (Fig. 4F; Supplemental Table S7). We found that *pmk-1* null and *pmk-1(CTEΔ)* mutants had a strong correlation in proteins enriched for K48 poly-ubiquitin linkages based on Pearson’s *r^2^* determination of variance (Fig. 4F). This finding paralleled their similar phenotypes in reduced longevity. In contrast, the *pmk-1* null mutant showed distinct enrichments of proteins tagged with K48-linked poly-ubiquitin compared to the cleavage-resistant *pmk-1(D327E)* mutant (Fig. 4F).

The K48-linked proteins were analyzed for GO term enrichments using the Wormbase enrichment toolkit. We found several cellular processes were inversely targeted for degradation in the *pmk-1(-)* null versus the cleavage-resistant *pmk-1(D327E)* mutants including cuticle structures, ATP hydrolysis, FAD binding, and chitin binding (Fig. 4G). Altogether, our findings suggest that the cleavage-resistant PMK-1 mutant behaves oppositely to the *pmk-1* null mutant and has heightened p38 signaling in both transcription and protein clearance programs. In addition to innate immunity, we find p38 signaling regulates both gene expression and degradation programs underlying tissue homeostasis.

### PMK-1 p38 MAPK regulates lysosome formation and activity

The lysosomal branch of protein clearance is key to support proteostasis. Long-lived mutants have recently been shown to retain young adult lysosome morphology as they progress into aging (Sun et al. 2020). Because we observed the altered lysosomal gene expression by loss of *pmk-1*, we further investigated if altered lysosomal function contributes to shortened lifespan of *pmk-1(-)* animal. Lysosomes function in complex organelle trafficking as they fuse with a variety of membrane-bound cargo. After lysosome fusion with the autophagosome, the autolysosome resolves cargo and subsequent tubulation regenerates lysosomes upon scission in a multi-step process. Tubules are thought to represent autolysosome extensions and can indicate a compromised ability to reform lysosomes as they progressively elongate (McGrath et al. 2021), consistent with failure to mitigate proteostasis challenges with aging.

Using the *P_CED-1_::nuc-1::*mCherry marker which localizes the NUC-1::mCherry fusion protein inside epidermal lysosomes (Sun et al. 2020), we monitored lysosome morphology during aging. For comparable quantitation, we imaged the same anatomical location across all animals and quantified all puncta and tubules in the views. At the onset of adulthood (day 1), lysosomes in wild-type animals were mostly particulate, with very few visible tubules (Fig. 5A). Following egg-laying (day 5) during early aging, wild-type animals displayed increased tubular lysosome structures (Fig. 5A-B). We observed that *pmk-1* null and CTEΔ mutants had very large lysosome structures at the onset of adulthood (day 1) (Fig. 5A-B; Supplemental Fig. S5A-B). The average lysosome particle size in *pmk-1* null mutants was doubled at day 1 of adulthood and continued with aging (day 5) (Fig. 5B). In contrast, cleavage-resistant *pmk-1(D327E)* mutants maintained small particle size and had much reduced tubular formation on day 5 of adulthood compared to wild-type (Fig. 5B). Long lived mutants such as *daf-2* null maintain very uniform particle size with few tubular structures (Sun et al. 2020) similar to our observation for *pmk-1(D327E)*. Intriguingly, wild-type and *ced-3* null mutants displayed comparable average particle size and tubular lysosome structures on day 1 and day 5 of adulthood (Fig. 5B). However, *ced-3* null mutants displayed increased variability in both particle and tubular structures. These results suggest that in addition to PMK-1, the caspase may act on additional factors that affect lysosome function.

**Figure 5.**
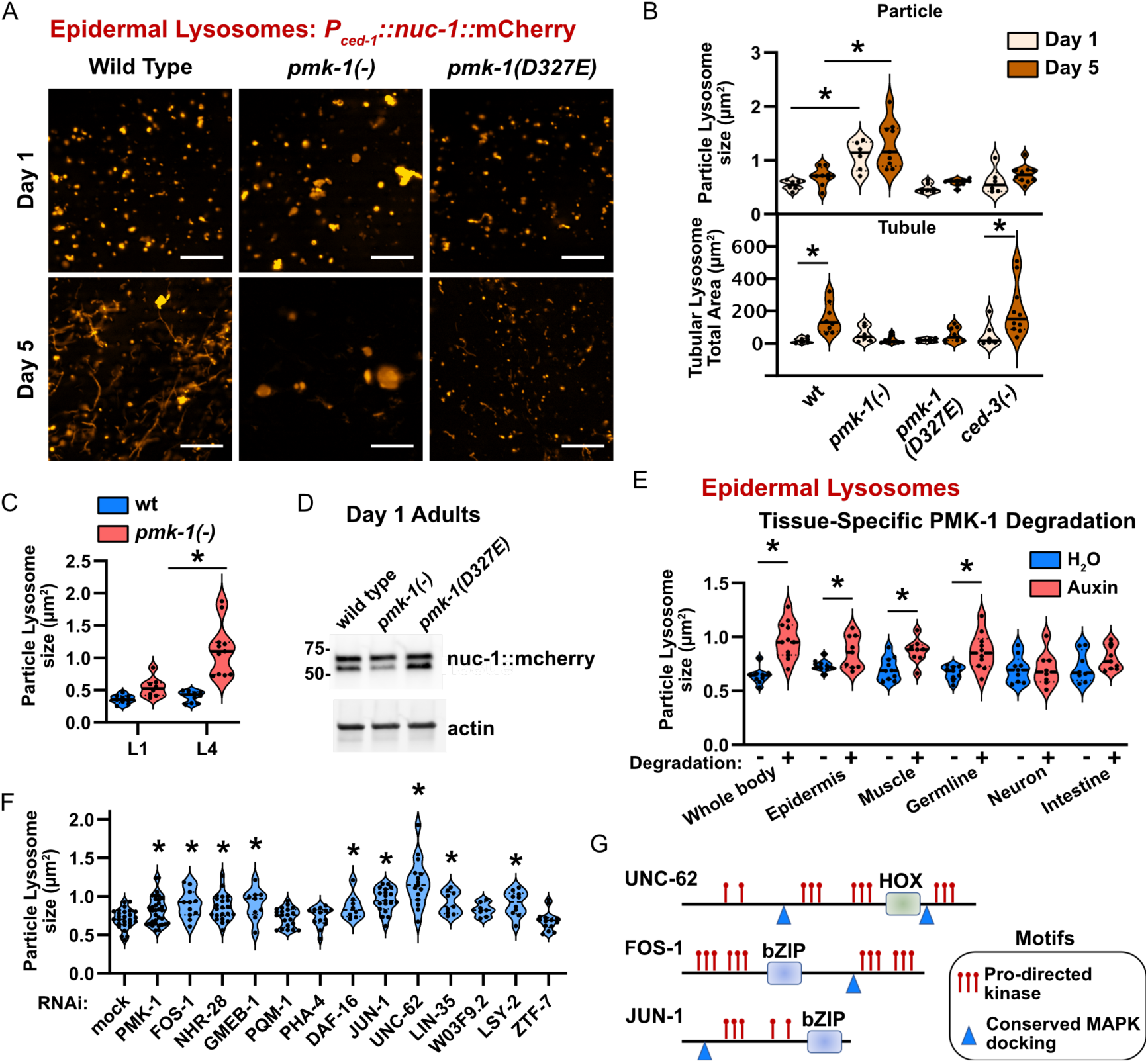
Cell autonomous and non-cell autonomous PMK-1 signaling is required for lysosome formation and activity in young adults. (*A-B*) Young adult epidermal lysosome structures monitored by DIC optics. Pseudo-colored epidermal lysosome marker (*P_CED-1_::NUC-1::mCherry*) during early aging (Day 1, and Day 5) and quantification. All particles and tubules in view were quantified and plotted as mean area (particles) or total area (tubules), +/- standard deviation. Each dot represents median value of all particles in one animal. Asterisk, *p < 0.05, Welch’s t Test* compared to wt. (*C*) First (L1) and final (L4) larval stage epidermal lysosome particle size quantification. (*D*) Western blot of NUC-1::mCherry protein showing *pmk-1-* dependent processing. (*E*) Test of cell autonomy for epidermal lysosome particle size. Day 1 adult quantification with tissue-specific elimination of *pmk-1.* (*F*) Functional test on epidermal lysosome particle size for TFs we identified by promoter analysis (Fig. 4). (*G*) Motif analysis for Proline-directed phosphorylation sites and conserved MAPK docking.

The lysosome enlargement for *pmk-1* null animals is progressive from the first (L1) to the final (L4) larval stages (Fig. 5C). To examine functional effects on lysosome proteolytic processing, we monitored NUC-1::mCherry processing for Day 1 adults and found that loss of *pmk-1* function had reduced processing (Fig. 5D). The findings of lysosomal morphology and function parallel well with our observation that *pmk-1* null mutants have shortened life spans whereas *pmk-1(D327E)* mutants display prolonged life spans.

We examined whether the defective enlargement of lysosomes was cell autonomous for PMK-1 signaling using a series of tissue-specific TIR-1 E3 ligase mutants that target the auxin-inducible degron on the N-terminus of PMK-1. Whole body, germline and gut TIR-1 strains were previously used (Zhang et al. 2015) and we generated muscle, epidermal, and neuronal TIR-1 strains (Supplemental Tables S1-S3; Materials and methods). Whole body elimination of PMK-1 function showed enlarged lysosomes at day 1 adulthood (Fig. 5E) similar to our findings for the *pmk-1* null mutant. Loss of PMK-1 specifically in epidermis also showed enlarged lysosomes (Fig. 5E) suggesting that lysosome trafficking has a cell-autonomous component. We further tested other tissues and revealed that muscle and germline PMK-1 functions also contribute to regulation of the epidermal lysosomal size indicating non-cell autonomous p38 signaling for lysosome formation. Because lysosomes are a major clearance pathway for proteostasis, our results indicate that epidermal proteostasis is supported directly within the epidermis as well as by distal tissues including the germline and muscle. Our findings for germline PMK-1 signaling supporting organismal proteostasis provided a potential mechanism for the observation that loss of *pmk-1* results in shortened lifespan when germline is left intact.

Based on our analysis for lysosomal enriched transcription factors (Fig. 4F), we tested the top 12 TFs for defects in lysosomal morphology. We found that the Meis homeobox ortholog UNC-62 had the most significant increase in epidermal lysosome particle size (Fig. 5F). Seven other transcription factors including FOS-1 (FOS), JUN-1 (JUN), NHR-28 (HNF4), GMEB-1 (GMEB), DAF-16 (FOXO), LIN-35 (RBL) and LSY-2 (ZNF18) also had significantly increased lysosome particle size (Fig. 5F), suggesting that PMK-1 works with a complex transcriptional network to regulate lysosomal genes. The p38 MAPK signaling phosphorylates serine or threonine residues on target proteins with a canonical proline-directed motif (S/TP) and substrates typically contain a MAPK-docking motif. UNC-62, FOS-1, and JUN-1 had enriched proline-directed motifs and each contained at least one docking motif (Fig. 5G; Supplemental Fig. S5C). Based on these structural motifs, it is conceivable that these transcription factors are directly regulated by a MAPK signaling cascade.

### PMK-1 p38 MAPK signaling supports neuronal integrity during aging

Lysosomes are critical to support organismal proteostasis. Neurons are particularly sensitive to declines in proteostasis. Loss of neuronal integrity is a major and significant feature of aging throughout animal phyla (Mattson and Magnus 2006; Toth et al. 2012). Because the gene expression data also revealed significant enrichments for neuronal genes regulated by PMK-1, we evaluated aging phenotypes associated with neurons in *C. elegans*. Using a pan-neuronal marker, *P_RGEF-1_::*DsRed (Kerk et al. 2017), we found that *pmk-1* null, *pmk-1(D327E),* and *ced-3* null mutants had normal neuronal morphology at day 1 young adulthood (Fig. 6A-B). By day 8 of adulthood, about half of wild-type animals had disrupted morphology in the large lateral mid-body sensory neurons with obvious puncta demonstrating a progressive decline in neuronal integrity with aging (Fig. 6C-D).

**Figure 6.**
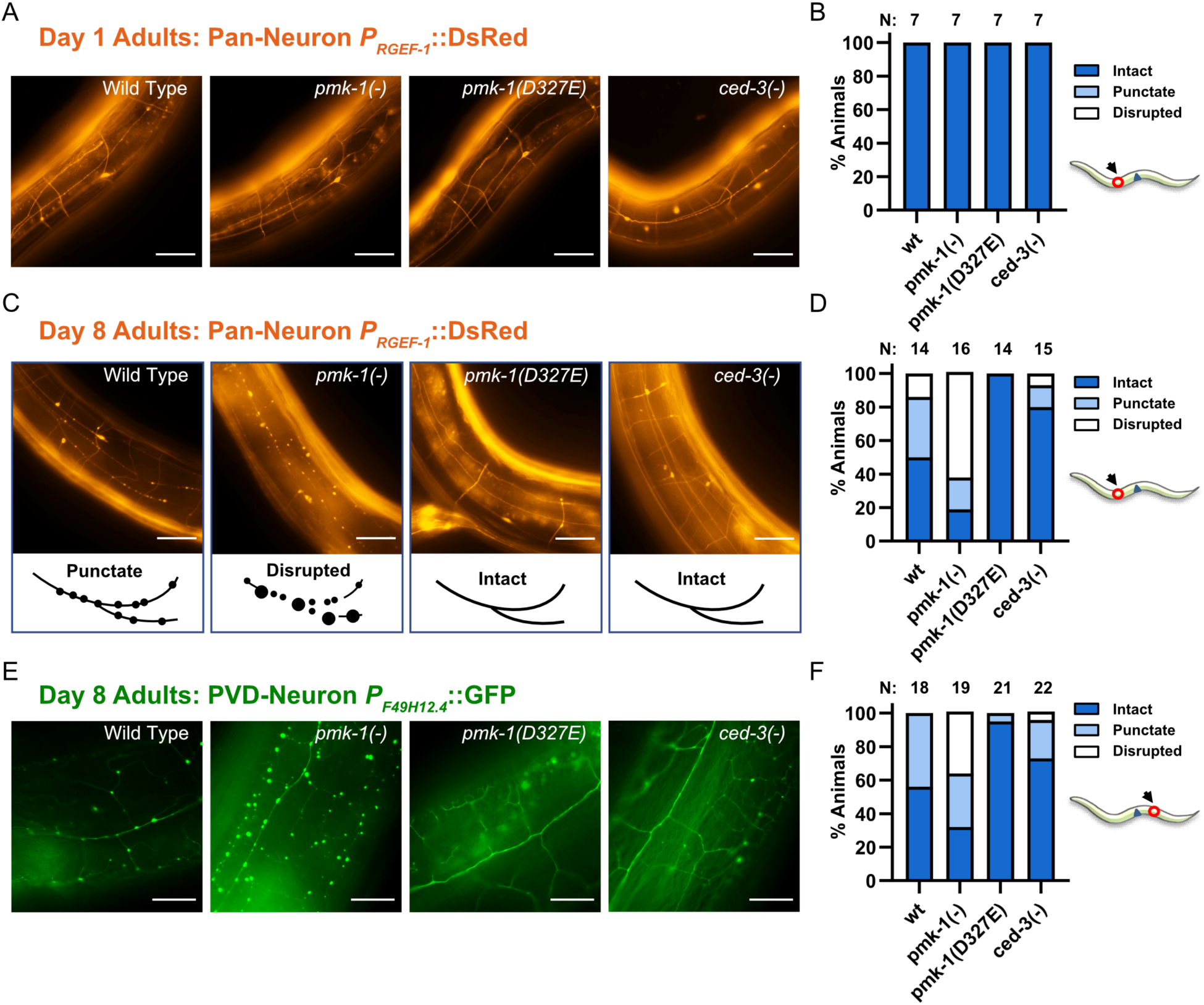
Caspase acting on PMK-1 regulates integrity of sensory neurons during aging. (*A-D*) Representative image and quantification of large lateral mid-body sensory neurons during aging depending on extent of PMK-1 signaling regulated by caspase cleavage. Neurons were visualized by DIC optics using a pan-neuronal marker (*P_RGEF-1_::DsRed,* pseudo color). Long continuous tracts without interruption (intact), thin nerves with punctate foci (punctate), and thin, disrupted nerves with large foci (disrupted). Number of animals indicated (N). Cartoon indicates position for imaging neuronal tracts. (*E-F*) Representative Images with pseudo-colored sensory neurons visualized by DIC optics using a PVD/AQR-specific marker (*F49H12.4::GFP*) and quantification. The PVD sensory neurons are anatomically associated with the epidermis.

Strikingly, this decline in neuronal integrity was accelerated in the majority of *pmk-1* null and CTEΔ mutants by day 8 of adulthood (Fig. 6C-D; Supplemental Fig. 5D-E). In contrast, we saw that both the cleavage-resistant *pmk-1(D327E)* and *ced-3* mutants were protective for neuronal morphology with aging (Fig. 6C-D). Using a PVD-specific reporter, we confirmed the PVD sensory neurons showed that loss of *pmk-1* function led to an accelerated decline in integrity with obvious puncta and lack of continuous tracts whereas the cleavage-resistant *pmk-1(D327E)* mutation showed a strong protective affect for neuronal integrity with few puncta and long continuous tracts and branching (Fig. 6E-F). These findings suggest that although p38 MAPK signaling is not required for development of these neurons it is a key regulator that can be modulated to protect neuronal integrity with aging. It is further intriguing that the PVD sensory neurons are anatomically interconnected with the epidermis. Disruption of p38 signaling in the epidermis compromised the epidermal lysosomes which may contribute to compromising the anatomically associated sensory neurons.

### Altering the non-phosphorylated PMK-1 pool modulates net p38 signaling to control growth and tissue dynamics

We observed the amount of phosphorylated PMK-1 is fairly constant during early aging but non-phosphorylated PMK-1 levels are dramatically downregulated and PMK-1 signaling is important during early aging to support longevity. These findings suggested the possibility that altering the phospho to non-phospho p38 ratio may impact net signaling. To test the effects of increasing the total PMK-1 pool, we used a constitutive promoter (*eft-3*) with permissive 3’ UTR element (*unc-54* 3’ UTR) to overexpress PMK-1. To limit PMK-1 overexpression and possible compensatory adaptation prior to experiments, we used an N-terminal auxin-induced degron (AID) tag, which did not affect the function of PMK-1 (Supplemental Fig. S2A). PMK-1 over-expression by this transgene was suppressed when treating animals with the auxin analog K-NAA (referred to as “auxin” throughout) (Fig. 7A).

**Figure 7.**
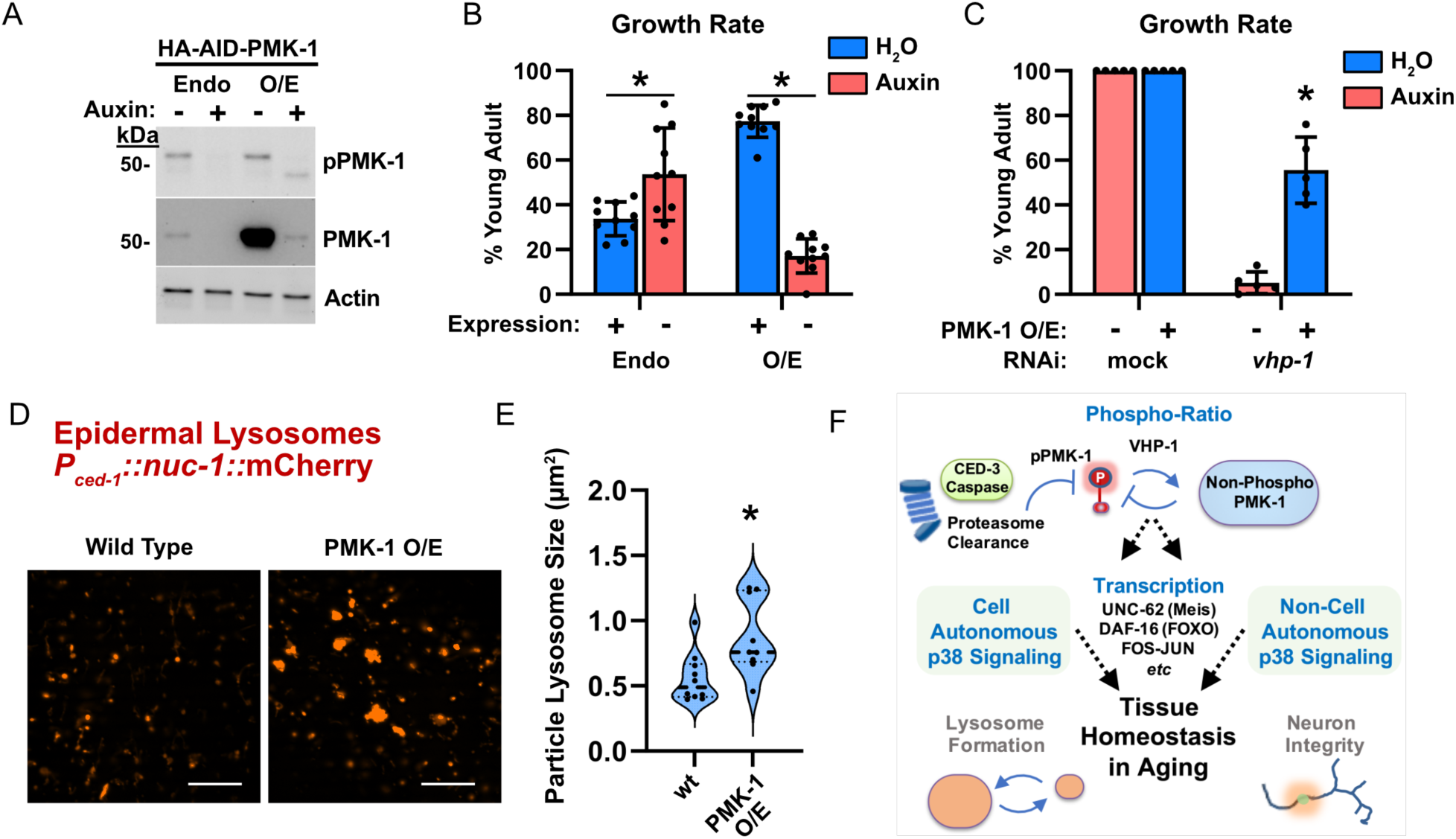
Increasing the non-phosphorylated PMK-1 pool protects against deleterious effects of PMK-1 hyperactivation. (*A*) Western blot of HA-AID-tagged PMK-1 in the endogenous locus (Endo) or an HA-AID-tagged PMK-1 transgene with *eft-3* promoter and *unc-54* 3’ UTR for overexpression (O/E). (*B-C*) Growth rate development assays to test if additional PMK-1 protein expression protects against PMK-1 hyperactivation. Synchronous larvae were allowed to develop for 60 hours on OP50 food (B) or HT115 mock and *vhp-1* RNAi (C). Animals develop faster on HT115 food than when fed OP50 food. Each dot corresponds to a plate of 25-40 animals. Error bars show standard deviation. (*D-E*) Day 1 adult epidermal lysosome structures monitored by DIC optics using epidermal lysosome marker (*P_CED-1_::NUC-1::mCherry*) and quantification. All particles in view were quantified and plotted as mean area +/- standard deviation. Each dot represents median values of all particles in one animal. Asterisk, *p < 0.05, Welch’s t Test* compared to wt. (*F*) Model showing the relative impact of phosphorylated and non-phosphorylated PMK-1 to net PMK-1 signaling integrated across tissues to support tissue homeostasis.

Overexpression of the wild-type PMK-1 resulted in about 40-fold increase in total PMK-1 but no increase in phosphorylated PMK-1 (Fig. 7A). This finding further supports our observation that phospho-PMK-1 levels are strictly limited in an animal. We previously showed that loss of *pmk-1* function results in faster larval development (Weaver et al. 2020). Consistent with this finding, knocking down endogenous PMK-1 protein with the auxin-induced degron (AID) system resulted in faster growth (Endo -, Fig. 7B). Interestingly, we also found that mutants with overexpressed PMK-1 phenocopied loss of *pmk-1* function with an enhanced growth rate (O/E +, Fig. 7B). These results were counterintuitive but suggested that overexpression of non-phosphorylated PMK-1 limits phospho-PMK-1 function and this is revealed by altering the stoichiometry.

There is no phospho-mimetic mutation for the TGY dual phospho-site able to mimic pTGpY. As such, we were unable to generate a suitable phospho-mimetic PMK-1 mutant. VHP-1 is a dual specificity phosphatase (DUSP) required to dephosphorylate PMK-1 (Kim et al. 2004; Mizuno et al. 2004). Therefore, we used *vhp-1(RNAi)* which will increase accumulation of phosphorylated PMK-1 (Supplemental Fig. 6A). Losing VHP-1 function during larval development by *vhp-1(RNAi)* causes a developmental delay for wild-type animals (Supplemental Fig. 6B) and this phenotype is PMK-1-dependent as seen by alleviation of stalled development in the absence of functional *pmk-1* (Supplemental Fig. 6B). The stalled development we observe in this system is consistent with previous findings by us and others when PMK-1 signaling is hyperactivated (Mizuno et al. 2004; Kim et al. 2016; Foster et al. 2020; Weaver et al. 2020). This treatment is therefore a useful tool to enhance phosphorylated PMK-1 levels to study PMK-1 signaling. To test the relationship of phospho-PMK-1 ratio to non-phospho-PMK-1 further, we examined rates of growth when altering both pPMK-1 and the total pool of PMK-1.

We observed that animals fed *vhp-1(RNAi)* without PMK-1 overexpression (auxin treated) had a marked developmental delay compared to mock (RNAi) (red bar, Fig. 7C). Interestingly, animals fed *vhp-1(RNAi)* but allowed to overexpress PMK-1 protein were significantly protected against the developmental delay with nearly half the animals reaching adulthood (blue bar, Fig. 7C). Moreover, we observed a similar result of overcoming the *vhp-1(RNAi)* delay by overexpressing a *pmk-1* mutant that cannot be phosphorylated (TGY mutated to AGF) (Supplemental Fig. 6C). These results suggest that increasing non-phosphorylated PMK-1 pool mitigated the effect of hyperactivated pPMK-1. Furthermore, when we overexpressed PMK-1, lysosome particle size was enlarged similar to *pmk-1* null (Fig. 7D-E). Altogether, our results suggest during normal aging, the net PMK-1 signaling output is modulated by altering the amount of non-phosphorylated PMK-1.

In total, our findings support a model whereby in addition to phosphorylation status, proteostasis mechanisms modulate PMK-1 function by regulating the balance of pPMK-1 and total PMK-1 pools. Moreover, PMK-1 has both cell autonomous and non-cell autonomous functions in supporting homeostasis by promoting lysosome function, an important clearance pathway for proteostasis and is critical to promote longevity (Fig. 7F).

## Discussion

Longevity requires the integration of many processes including resolving protein aggregates, nutrient sensing, and genomic integrity among other factors (Riera et al. 2016). Succumbing to pathogen infection is a common feature of animal aging. PMK-1 was previously shown to activate the majority of pathogen response genes via the transcription factor ATF-7, supporting survival on pathogenic *Pseudomonas* bacteria (Fletcher et al. 2019). Additionally, regulation of the immunometabolic program by the PMK-1-ATF-7 circuit is a key determinant for longevity in response to nutrient availability or infection (Wu et al. 2019). These functions are critical for survival when faced with fluctuating environmental conditions.

Here we demonstrate an under-recognized function of a p38 family member in aging apart from its stress sensing functions supporting adult survival with pathogen insult. Although loss of *pmk-1* function does not compromise development and even accelerates developmental speed, loss of the gene results in significantly compromised neuronal integrity and lysosomal structures in adulthood resulting in shortened lifespan in the presence of the germline. The *pmk-1(-)* null mutant treated with FUDR to eliminate germline has been shown to have normal life span in the absence of infection (Troemel et al. 2006; Wu et al. 2019). Altogether these results are consistent with the idea that the proteostasis load of germline proliferation impact longevity (Labbadia and Morimoto 2015; Hipp et al. 2019). This is further supported by the observation that eliminating p38 signaling in the germline compromised lysosome structure in hypodermis.

From our findings, we raised the question of how overexpression of PMK-1 protein phenocopies the *pmk-1* null mutation. It is feasible that innate regulatory mechanisms could exploit distinct protein-protein interactions between the phosphorylated and non-phosphorylated forms of PMK-1. A previous model predicted that MAPK interactions with scaffold proteins may ultimately regulate net signaling output from MAPKs (Levchenko et al. 2000). This idea is consistent with the observation that *pmk-1(D327E)* behaves as a gain-of-function mutation despite accumulating less total protein. It is even possible this mechanism is operative during aging thereby enhancing p38 signaling. By down regulating non-phospho-PMK-1 while maintaining the absolute amount of phospho-PMK-1 with age, the output of p38 signaling is increased for the maintenance of tissue homeostasis without the hyper-activation of stress-responses.

The function of proteostasis in mitigating stress responses such as unfolded protein responses have clear consequences on longevity and degenerative diseases (Morley and Morimoto 2004; Taylor and Dillin 2013; Morimoto 2020). Recent findings have illuminated roles for p38 MAPK in coordinating stress signaling within cells including modulating ER stress resistance by cell surface hyaluronidase TMEM2 independent of canonical UPR (Schinzel et al. 2019) and PERK-dependent recruitment of MKK4 and p38 targeting LAMP2A on lysosomes to activate chaperone-mediated autophagy (Li et al. 2017). Lysosomes have recently been identified in a fat-to-neuron lipid-signaling function in *C. elegans* by mobilizing a polyunsaturated fatty acid with lipolysis then subsequent transport by a lipid binding protein to activate the nuclear receptor NHR-49, and activation of neuropeptide signaling to promote longevity (Savini et al. 2022). Our current study unmasked p38 MAPK signaling working with a complex network of transcriptional factors to maintain lysosome function during aging in the absence of stress. Altogether, these findings underscore the importance of proteostasis in longevity.

Following injury, CED-3 was previously shown to promote neuronal regeneration in *C. elegans* (Pinan-Lucarre et al. 2012) suggesting a caspase may have multiple targets in neuronal protection. Further, the observation that loss of CED-3 caspase did not always correspond to cleavage-resistant PMK-1 phenotypes underscores the complexity of the cross-talk between the caspase and MAPK signaling pathways. Future studies will certainly prove out the intricacies of caspases working to sculpt multiple aspects of signal transduction in these and other functions.

Recent findings have revealed a complex relationship for p38 signaling in models of neuronal degeneration. Neurodegeneration as a result of expressing mutant TDP-43 or FUS is driven by an innate immune response dependent on PMK-1 signaling (Veriepe et al. 2015). In contrast, loss of mitochondria resulting in neurodegeneration is alleviated by activating CaMKII (UNC-43) signaling leading to PMK-3 activation and ultimately CEBP (Ding et al. 2022). Intriguingly, this rescue is specific to PMK-3 signaling and not PMK-1 signaling (Ding et al. 2022). In the current study, we identified *pmk-1-*dependent signaling for key gene expression programs to support sensory neuronal integrity during aging in the absence of a stressor. Because we have observed that PMK-1 signaling has both cell-autonomous and non-cell-autonomous components and sensory neurons are intimately connected with epidermis, it is possible that the neuronal effect we observe is a consequence of diminished signaling in surrounding tissue. Future studies are warranted to investigate this further.

Altogether, our results suggest that moderate enhancement of p38 signaling during aging may have unexpected protective impacts on tissue homeostatic processes by strengthening proteostasis and protein clearing mechanisms. The enhanced p38 signaling during aging is achieved by proteostatic regulation that decreases the amount of total p38 protein without affecting the amount of phosphorylated-p38, therefore preventing hyperactivation of immune or stress response.

## Materials and Methods

### Protein modeling

AlphaFold (Jumper et al. 2021) was used to generate a model of *C. elegans* PMK-1 using p38α (PDB code 2OKR) and p38γ (PDB code 6UNA). Model confidence around the caspase cleavage site is very high (pLDDT > 90).

### *In vitro* cleavage assay

Caspase cleavage reactions were performed as previously reported (Weaver et al. 2017). *C. elegans* PMK-1 and *Drosophila* p38b were both cleaved with *Sf21* insect cell extract containing active caspase. Purified human caspases-2, -3, and -8 were incubated with human p38α and p38γ. 35S-labeled substrates were synthesized fresh before cleavage reactions. Cleavage reactions were stopped by adding sample buffer and heating to 85°C. Samples were resolved by SDS-PAGE on 4-16% gradient gels and dried before imaging.

### Strains used in this study and culturing conditions

For list of strains used in this study, see Table S1. Standard *C. elegans* culture conditions and RNAi treatment were the same as previously described (Weaver et al. 2020). In brief, NGM agar plates with OP50 lawn were used for standard maintenance, growth and aging assay. HT115 culture was seeded on NGM agar plates containing 1 mM IPTG and 200 μg/mL ampicillin for RNAi treatments and assays.

### CRISPR-Cas9 mutagenesis

For list of all sgRNAs and repair templates used in this study, see Supplemental Tables S2 and S3. Endogenous HA-AID-GS and HA-GS tagged PMK-1 or HA-GS tagged PMK-1 (D327E) mutant were generated by CRISPR/Cas9. In brief the rescue DNA was assembled using NEB HiFi DNA assemble Kit and purified PCR product was used as the rescue template. The AID degron tag was amplified from pLZ31 (Zhang et al. 2015) with modification. The injection constructs mix containing the sgRNA target the 1st exon of pmk-1 (pDD162_C0025, Key Resources Table) were injected into N2 strain or pmk-1 (D327E) mutant (Weaver et al. 2020) respectively. To generate the overexpression mutant, a single copy of *ha-aid-gs-pmk-1, ha-aid-gs-pmk-1 (D327E)* or *ha-aid-gs-pmk-1 (T191A, Y193F)* flanked by the *eft-3* promoter and *unc-54* 3’ UTR were inserted into Mos I site oxTi365 on Chr V by CRISPR/Cas9 following the SEC methods (Dickinson et al. 2015). The overexpression mutants were generated by injecting constructs into the CA1200 strain (Zhang et al. 2015).

### Auxin-induced degron (AID) system culture condition

To conditionally inactivate PMK-1 or over-expressed PMK-1, we used the Auxin-Induced Degron (AID) system. The auxin analog K-NAA was used in lieu of auxin for this study. For culture conditions, we sterile-filtered the auxin analog K-NAA dissolved in MilliQ H2O and added to standard NGM agar to a final concentration of 0.1 mM K-NAA followed by seeding of OP50 as normal. We find that adults maintained on K-NAA plates transferred to standard NGM/OP50 media begin to restore PMK-1 expression within 24 hours and reach wild-type levels within another two days of adulthood.

For tissue specific PMK-1 degradation, endogenous PMK-1 was tagged with HA-AID at N-terminus using CRSIPR-Cas9 mutagenesis and crossed into strains carrying TIR1 E3 ligase expressed under different promoters including: whole body (*eft-3,* CA1200), Intestine (*ges-1,* CA1209) and germ line (*sun-1,* CA1199) as well as additional strains generated in this study including: epidermis (*col-10)*, body wall muscle (*myo-3),* pan-neuron (*rgef-1)*.

### Growth rate assay

Five gravid young adults were allowed to lay eggs for 1 hour on each plate and removed. Animals were staged at the indicated time intervals. Prior to testing, animals were maintained under stress-free conditions at 20°C for multiple generations, including no starvation and no obvious contaminations.

### Aging assay

Mid-L4 stage animals were placed on OP50 NGM (no FUDR) plates with 20 animals per plate and 5 plates per genotype. The next day animals were confirmed to be egg-laying and was defined as the Day 1 of the aging assay. Worms were moved to a fresh NGM plates daily for the first five days and subsequently were moved every 4-5 days.

### Imaging

The epidermal lysosomal marker (*P_CED-1_::nuc-1::*mCherry) (Sun et al. 2020) was used to image epidermal lysosome structure during aging. The pan-neuronal marker (*P_RGEF-1_::*DsRed) (Kerk et al. 2017) and the PVD/ AQR-neuronal marker (*P_F49H12.4_::*GFP) was used to image neurons. Zeiss AxioImager M2 with a black and white Hamamatsu ORCA C13440 camera was used for DIC images. All images were taken at the same magnification for the same amount of time for comparisons as indicated in the relevant figure legends.

### Western blots

Synchronous L1 stage animals were obtained following standard alkaline bleach to obtain eggs as previously described (Weaver et al. 2017). L2, L3, L4, and day 1 adults were collected at 8, 18, 30, and 48 hours post-feeding. Animals were washed off plates and snap frozen with liquid nitrogen. Pellets were sonicated in lysis buffer containing 10 mM Tris pH 7.4, 1 mM EDTA, 150 mM NaCl, 0.5 % NP-40, Halt™ Protease and Phosphatase Inhibitor Cocktails (Fisher Scientific, PI78440) and kept cool using ice. Approximately 4 – 6 µg of total protein was loaded per well and resolved on 4 -20% gradient acrylamide gels. Anti-HA antibody from Rabbit (Cell Signaling Technology, 3724S) or from Mouse (Cell Signaling Technology, 2367S) were used at 1:1000 dilution. Anti Phospho-p38 (Cell Signaling Technology, 4511S) was used at 1:2000 dilution. Anti-α-tubulin (Sigma-Aldrich, T5168) or Anti-Actin (Bio-Rad, 12004163) antibodies were used at 1:4000 and 1:2000 respectively for loading controls as indicated.

### *In vitro* stability assay

TNT Rabbit reticulocytes were used to radiolabel proteins with 35S-Methionine for 1 hour at 30°C. Products were then diluted 1:5 into 25 mM Tris, pH 8.0, 0.25 mM EDTA, 0.25 mM sucrose, and 2.5% glycerol. Mock reactions were collected directly into 3 volumes of 2X SDS buffer with 5% β-mercaptoethanol. Two additional reactions were incubated at either 20°C or 37°C for 48 hours and then quenched with 3 volumes of 2X SDS buffer. One-fourth of samples were resolved on 4 -20% acrylamide gels, fixed with 50% methanol and 10% glacial acetic acid, protected with 10% glycerol, and dried onto Whatman filter paper. Dried gels were exposed to phospho-imager screens over two nights and imaged on phospho-imager.

### Proteasome, lysosome and autophagy inhibition assays

Synchronous L1 stage animals were fed OP50 for 40hrs and washed off plates into S-basal containing Bortezomib (130µM), Bafliomycin-A1 (50µM), 3-MA(50mM) or DMSO (0.1%) with OP50 food. The liquid culture was incubated at 20°C for 6 hrs with 250rpm shaking. Then the worms were washed 4 times with M9 and the pellet was frozen in liquid N2 and stored at -80°C until Western Blot analysis.

### Statistics

Statistics were reported in the Legends with *p* values indicated throughout.

### mRNA-Seq and transcription factor binding site analyses

Synchronous young adult stage animals were collected into TRIzol™ Reagent (Invitrogen, 15596-026). Total RNA was extracted. Three replicates each of wild-type, *pmk-1(-)*, and *pmk-1(D327E)* mutants were collected. For *daf-16* mutants, *daf-16(-);pmk-1(-)* and *daf-16(D327E)* mutants, two replicates each for *m26* and *mgDf50* alleles were used. Libraries were generated and sequenced and analyzed for differential gene expression by the UTSW McDermott Center Next Generation sequencing core. Wormbase gene set enrichment analysis tools were used for gene ontology analyses. Transcription factor binding sites analysis was done using modENCODE (from Wormbase JBrowse) database. All the TFs binds to ∼2kb upstream of the transcription start site were recorded and data from all developmental stages were pooled (Supplemental Table S6).

### Reverse transcription and quantitative PCR

Total RNA was extracted using TRIzol™ Reagent (Invitrogen, 15596-026) from animals with different genotypes and developmental stages. For cDNA synthesis, 1µg of total RNA was used as input for all samples. The iScript RT Supermix for RT-qPCR (BioRad, 1708841) was used for cDNA synthesis. The CFX Realtime PCR system (BioRad) with iTaq Univeral SYBR Green Supermix (BioRad, 1725122) was used for qPCR analysis. Each condition was tested with 3 biological replicates. *C. elegans ama-1* was used as the reference gene for normalization in all experiments.

### PTM Quantification with Mass Spectrometry

Synchronized young adult animals expressing N-terminal HA-tagged PMK-1 were used for immunoprecipitation with anti-HA-Tag (C29F4) rabbit mAb (Cell Signaling Technology 11846) and eluted with 2x laemmli buffer. Eluted protein was resolved by SDS-PAGE and gel slices of PMK-1 band were sent for protein ID using Mass Spectrometry. The ratio of the abundance of T196 phospho-peptide to the unphosphorylated peptide was used to estimate the percentage of phosphorylated PMK-1.

### Lys48 linkage Ub Pull Down and Mass Spectrometry

Synchronized young adult animals were treated with Bortezomib (60µM) for 8hrs and worm pellets were washed 4 times and flash frozen in liquid nitrogen. Lys48-specific anti-Ubiquitin rabbit clone Apu2 (Sigma 05-1207 and ZRB2150) was used for immunoprecipitation at 4°C for 4 hours. The immune-complex was captured using Pierce MS-Compatible Magnetic IP Kit Protein A/G (ThermoFisher 90409) and eluted with 2x laemmli buffer. Eluted protein was resolved by SDS-PAGE and gel slices above 50KDa were sent for protein ID using mass spectrometry. Mass-spec and protein identification were performed by the UTSW proteomics core.

## Data and software availability

All mRNA-Seq data are deposited in GEO accession GSE192941

## Acknowledgements

We thank David Mangelsdorf, Leon Avery, James Collins, Michael Reese, and David Corey for helpful discussions, Matthew Sieber for helpful discussions and providing *Drosophila* mRNA, the CGC (funded by NIH Office of Research Infrastructure Programs [P40 OD010440]) for materials; WormBase and UniProt databases. RNA-seq was performed by the McDermott Center Sequencing Core and data analysis was provided by the McDermott Center Bioinformatics Lab. Mass Spectrometry and data analysis was provided by UTSW Proteomics Core. This work is supported by Welch Foundation grants I-2022-20190330 (BPW) and I1243 (MHC), and National Institutes of Health grant R35GM133755 (BPW). The funders had no role in study design, data acquisition, decision to publish, or preparation of the manuscript.

## Author Contributions

W.Y., Y.M.W., M.H.C, and B.P.W. conceived the study and designed research; W.Y., Y.M.W., S.E., C.A.T., and B.P.W. performed research; W.Y., Y.M.W., S.E., C.A.T., M.H.C, and B.P.W. analyzed data; W.Y., Y.M.W., M.H.C and B.P.W wrote the manuscript; M.H.C. and B.P.W. acquired funding.

## References

1. Arthurton L, Nahotko DA, Alonso J, Wendler F, Baena-Lopez LA. 2020. Non-apoptotic caspase activation preserves Drosophila intestinal progenitor cells in quiescence. EMBO Rep 21: e48892.

2. Bell RAV, Al-Khalaf MH, Brunette S, Alsowaida D, Chu A, Bandukwala H, Dechant G, Apostolova G, Dilworth FJ, Megeney LA. 2022. Chromatin Reorganization during Myoblast Differentiation Involves the Caspase-Dependent Removal of SATB2. Cells 11.

3. Ben-Zvi A, Miller EA, Morimoto RI. 2009. Collapse of proteostasis represents an early molecular event in Caenorhabditis elegans aging. Proc Natl Acad Sci U S A 106: 14914–14919.

4. Bluher M, Kahn BB, Kahn CR. 2003. Extended longevity in mice lacking the insulin receptor in adipose tissue. Science 299: 572–574.

5. Canovas B, Nebreda AR. 2021. Diversity and versatility of p38 kinase signalling in health and disease. Nat Rev Mol Cell Biol 22: 346–366.

6. Clancy DJ, Gems D, Harshman LG, Oldham S, Stocker H, Hafen E, Leevers SJ, Partridge L. 2001. Extension of life-span by loss of CHICO, a Drosophila insulin receptor substrate protein. Science 292: 104–106.

7. Dick SA, Chang NC, Dumont NA, Bell RA, Putinski C, Kawabe Y, Litchfield DW, Rudnicki MA, Megeney LA. 2015. Caspase 3 cleavage of Pax7 inhibits self-renewal of satellite cells. Proc Natl Acad Sci U S A 112: E5246–E5252.

8. Dickinson DJ, Pani AM, Heppert JK, Higgins CD, Goldstein B. 2015. Streamlined Genome Engineering with a Self-Excising Drug Selection Cassette. Genetics 200: 1035–1049.

9. Ding C, Wu Y, Dabas H, Hammarlund M. 2022. Activation of the CaMKII-Sarm1-ASK1-p38 MAP kinase pathway protects against axon degeneration caused by loss of mitochondria. Elife 11.

10. Fletcher M, Tillman EJ, Butty VL, Levine SS, Kim DH. 2019. Global transcriptional regulation of innate immunity by ATF-7 in C. elegans. PLoS Genet 15: e1007830.

11. Foster KJ, Cheesman HK, Liu P, Peterson ND, Anderson SM, Pukkila-Worley R. 2020. Innate Immunity in the C. elegans Intestine Is Programmed by a Neuronal Regulator of AWC Olfactory Neuron Development. Cell Rep 31: 107478.

12. Hipp MS, Kasturi P, Hartl FU. 2019. The proteostasis network and its decline in ageing. Nat Rev Mol Cell Biol 20: 421–435.

13. Holzenberger M, Dupont J, Ducos B, Leneuve P, Geloen A, Even PC, Cervera P, Le Bouc Y. 2003. IGF-1 receptor regulates lifespan and resistance to oxidative stress in mice. Nature 421: 182–187.

14. Inoue H, Hisamoto N, An JH, Oliveira RP, Nishida E, Blackwell TK, Matsumoto K. 2005. The C. elegans p38 MAPK pathway regulates nuclear localization of the transcription factor SKN-1 in oxidative stress response. Genes Dev 19: 2278–2283.

15. Jumper J, Evans R, Pritzel A, Green T, Figurnov M, Ronneberger O, Tunyasuvunakool K, Bates R, Zidek A, Potapenko A et al. 2021. Highly accurate protein structure prediction with AlphaFold. Nature 596: 583–589.

16. Kerk SY, Kratsios P, Hart M, Mourao R, Hobert O. 2017. Diversification of C. elegans Motor Neuron Identity via Selective Effector Gene Repression. Neuron 93: 80–98.

17. Kim DH, Liberati NT, Mizuno T, Inoue H, Hisamoto N, Matsumoto K, Ausubel FM. 2004. Integration of Caenorhabditis elegans MAPK pathways mediating immunity and stress resistance by MEK-1 MAPK kinase and VHP-1 MAPK phosphatase. Proc Natl Acad Sci U S A 101: 10990–10994.

18. Kim KW, Thakur N, Piggott CA, Omi S, Polanowska J, Jin Y, Pujol N. 2016. Coordinated inhibition of C/EBP by Tribbles in multiple tissues is essential for Caenorhabditis elegans development. BMC Biol 14: 104.

19. Labbadia J, Morimoto RI. 2015. The biology of proteostasis in aging and disease. Annu Rev Biochem 84: 435–464.

20. Levchenko A, Bruck J, Sternberg PW. 2000. Scaffold proteins may biphasically affect the levels of mitogen-activated protein kinase signaling and reduce its threshold properties. Proc Natl Acad Sci U S A 97: 5818–5823.

21. Li W, Zhu J, Dou J, She H, Tao K, Xu H, Yang Q, Mao Z. 2017. Phosphorylation of LAMP2A by p38 MAPK couples ER stress to chaperone-mediated autophagy. Nat Commun 8: 1763.

22. Lin K, Dorman JB, Rodan A, Kenyon C. 1997. daf-16: An HNF-3/forkhead family member that can function to double the life-span of Caenorhabditis elegans. Science 278: 1319–1322.

23. Mattson MP, Magnus T. 2006. Ageing and neuronal vulnerability. Nat Rev Neurosci 7: 278–294.

24. McGrath MJ, Eramo MJ, Gurung R, Sriratana A, Gehrig SM, Lynch GS, Lourdes SR, Koentgen F, Feeney SJ, Lazarou M et al. 2021. Defective lysosome reformation during autophagy causes skeletal muscle disease. J Clin Invest 131.

25. Mishra N, Wei H, Conradt B. 2018. Caenorhabditis elegans ced-3 Caspase Is Required for Asymmetric Divisions That Generate Cells Programmed To Die. Genetics 210: 983–998.

26. Mizuno T, Hisamoto N, Terada T, Kondo T, Adachi M, Nishida E, Kim DH, Ausubel FM, Matsumoto K. 2004. The Caenorhabditis elegans MAPK phosphatase VHP-1 mediates a novel JNK-like signaling pathway in stress response. EMBO J 23: 2226–2234.

27. Morimoto RI. 2020. Cell-Nonautonomous Regulation of Proteostasis in Aging and Disease. Cold Spring Harb Perspect Biol 12.

28. Morley JF, Morimoto RI. 2004. Regulation of longevity in Caenorhabditis elegans by heat shock factor and molecular chaperones. Mol Biol Cell 15: 657–664.

29. Ogg S, Paradis S, Gottlieb S, Patterson GI, Lee L, Tissenbaum HA, Ruvkun G. 1997. The Fork head transcription factor DAF-16 transduces insulin-like metabolic and longevity signals in C. elegans. Nature 389: 994–999.

30. Peterson ND, Icso JD, Salisbury JE, Rodriguez T, Thompson PR, Pukkila-Worley R. 2022. Pathogen infection and cholesterol deficiency activate the C. elegans p38 immune pathway through a TIR-1/SARM1 phase transition. Elife 11.

31. Pinan-Lucarre B, Gabel CV, Reina CP, Hulme SE, Shevkoplyas SS, Slone RD, Xue J, Qiao Y, Weisberg S, Roodhouse K et al. 2012. The core apoptotic executioner proteins CED-3 and CED-4 promote initiation of neuronal regeneration in Caenorhabditis elegans. PLoS Biol 10: e1001331.

32. Raman M, Chen W, Cobb MH. 2007. Differential regulation and properties of MAPKs. Oncogene 26: 3100–3112.

33. Riera CE, Merkwirth C, De Magalhaes Filho CD, Dillin A. 2016. Signaling Networks Determining Life Span. Annu Rev Biochem 85: 35–64.

34. Savini M, Folick A, Lee YT, Jin F, Cuevas A, Tillman MC, Duffy JD, Zhao Q, Neve IA, Hu PW et al. 2022. Lysosome lipid signalling from the periphery to neurons regulates longevity. Nat Cell Biol 24: 906–916.

35. Schinzel RT, Higuchi-Sanabria R, Shalem O, Moehle EA, Webster BM, Joe L, Bar-Ziv R, Frankino PA, Durieux J, Pender C et al. 2019. The Hyaluronidase, TMEM2, Promotes ER Homeostasis and Longevity Independent of the UPR(ER). Cell 179: 1306–1318 e1318.

36. Sun Y, Li M, Zhao D, Li X, Yang C, Wang X. 2020. Lysosome activity is modulated by multiple longevity pathways and is important for lifespan extension in C. elegans. Elife 9.

37. Tatar M, Kopelman A, Epstein D, Tu MP, Yin CM, Garofalo RS. 2001. A mutant Drosophila insulin receptor homolog that extends life-span and impairs neuroendocrine function. Science 292: 107–110.

38. Taylor RC, Dillin A. 2013. XBP-1 is a cell-nonautonomous regulator of stress resistance and longevity. Cell 153: 1435–1447.

39. Toth ML, Melentijevic I, Shah L, Bhatia A, Lu K, Talwar A, Naji H, Ibanez-Ventoso C, Ghose P, Jevince A et al. 2012. Neurite sprouting and synapse deterioration in the aging Caenorhabditis elegans nervous system. J Neurosci 32: 8778–8790.

40. Troemel ER, Chu SW, Reinke V, Lee SS, Ausubel FM, Kim DH. 2006. p38 MAPK regulates expression of immune response genes and contributes to longevity in C. elegans. PLoS Genet 2: e183.

41. Tullet JM, Hertweck M, An JH, Baker J, Hwang JY, Liu S, Oliveira RP, Baumeister R, Blackwell TK. 2008. Direct inhibition of the longevity-promoting factor SKN-1 by insulin-like signaling in C. elegans. Cell 132: 1025–1038.

42. Veriepe J, Fossouo L, Parker JA. 2015. Neurodegeneration in C. elegans models of ALS requires TIR-1/Sarm1 immune pathway activation in neurons. Nat Commun 6: 7319.

43. Visscher M, De Henau S, Wildschut MHE, van Es RM, Dhondt I, Michels H, Kemmeren P, Nollen EA, Braeckman BP, Burgering BMT et al. 2016. Proteome-wide Changes in Protein Turnover Rates in C. elegans Models of Longevity and Age-Related Disease. Cell Rep 16: 3041–3051.

44. Walther DM, Kasturi P, Zheng M, Pinkert S, Vecchi G, Ciryam P, Morimoto RI, Dobson CM, Vendruscolo M, Mann M et al. 2015. Widespread Proteome Remodeling and Aggregation in Aging C. elegans. Cell 161: 919–932.

45. Weaver BP, Weaver YM, Mitani S, Han M. 2017. Coupled Caspase and N-End Rule Ligase Activities Allow Recognition and Degradation of Pluripotency Factor LIN-28 during Non-Apoptotic Development. Dev Cell 41: 665–673.

46. Weaver BP, Weaver YM, Omi S, Yuan W, Ewbank JJ, Han M. 2020. Non-Canonical Caspase Activity Antagonizes p38 MAPK Stress-Priming Function to Support Development. Dev Cell 53: 358–369 e356.

47. Weaver BP, Zabinsky R, Weaver YM, Lee ES, Xue D, Han M. 2014. CED-3 caspase acts with miRNAs to regulate non-apoptotic gene expression dynamics for robust development in C. elegans. Elife 3: e04265.

48. Wu Z, Isik M, Moroz N, Steinbaugh MJ, Zhang P, Blackwell TK. 2019. Dietary Restriction Extends Lifespan through Metabolic Regulation of Innate Immunity. Cell Metab 29: 1192–1205 e1198.

49. Youngman MJ, Rogers ZN, Kim DH. 2011. A decline in p38 MAPK signaling underlies immunosenescence in Caenorhabditis elegans. PLoS Genet 7: e1002082.

50. Zhang L, Ward JD, Cheng Z, Dernburg AF. 2015. The auxin-inducible degradation (AID) system enables versatile conditional protein depletion in C. elegans. Development 142: 4374–4384.

