## Supplemental Figures for "Balancing p38 MAPK Signaling with Proteostasis Mechanisms Supports Tissue Integrity during Aging in *C. elegans*"

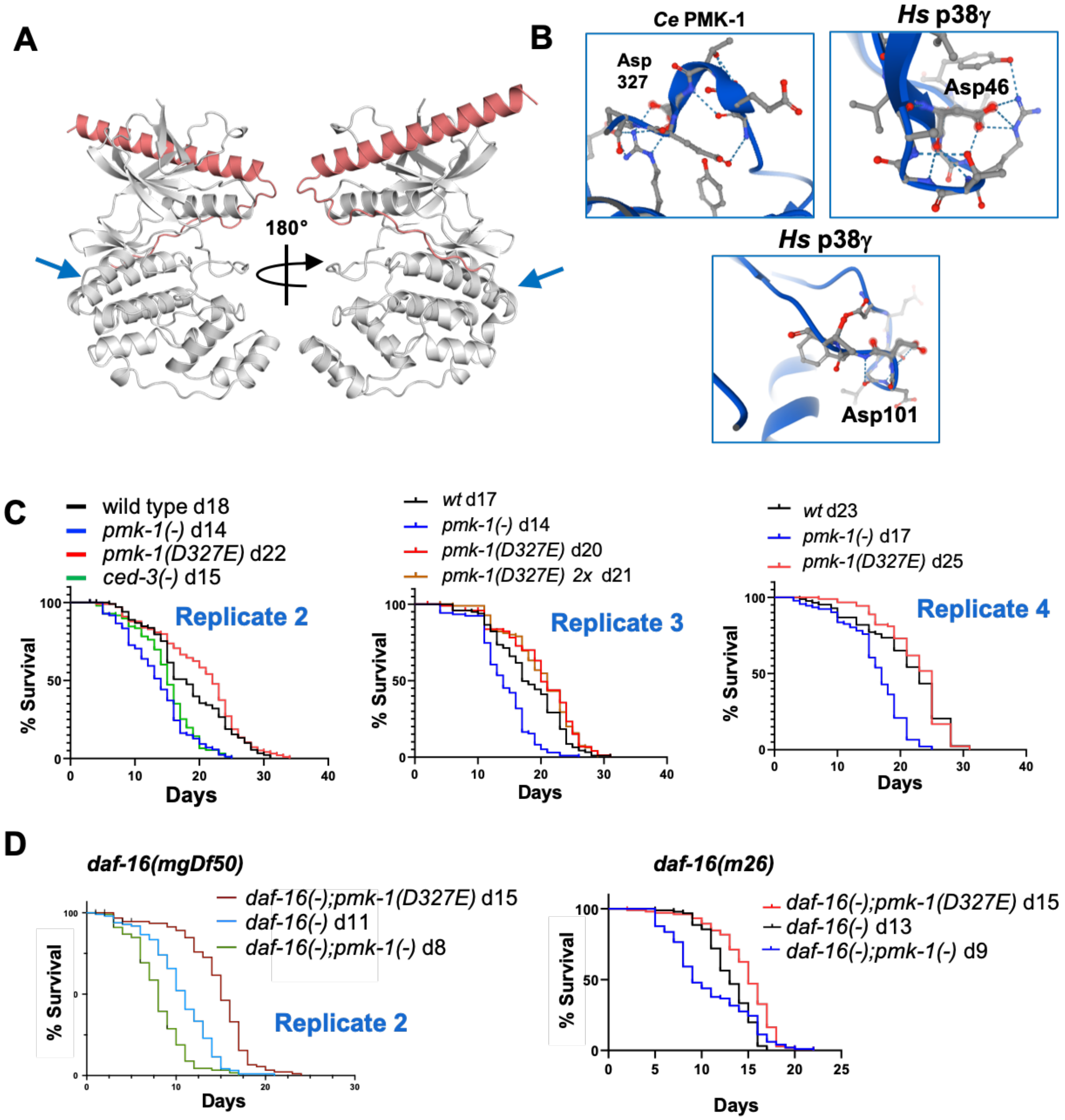


**Suppl. Fig. S1.** Replicates for cleavage resistant PMK-1 extends lifespan

**A-B)** Alphafold model of PMK-1. Arrows indicate cleavage site. Cleavage sites are solvent accessible in loops.

**C)** Replicates of *pmk-1* null and *pmk-1(D327E)* mutants with and without outcross in aging.

**D)** Replicates of effects of *pmk-1* null and *pmk-1(D327E)* mutations on *daf-16* mutant aging. For all aging assay replicates and statistics (Supplementary Table S4).

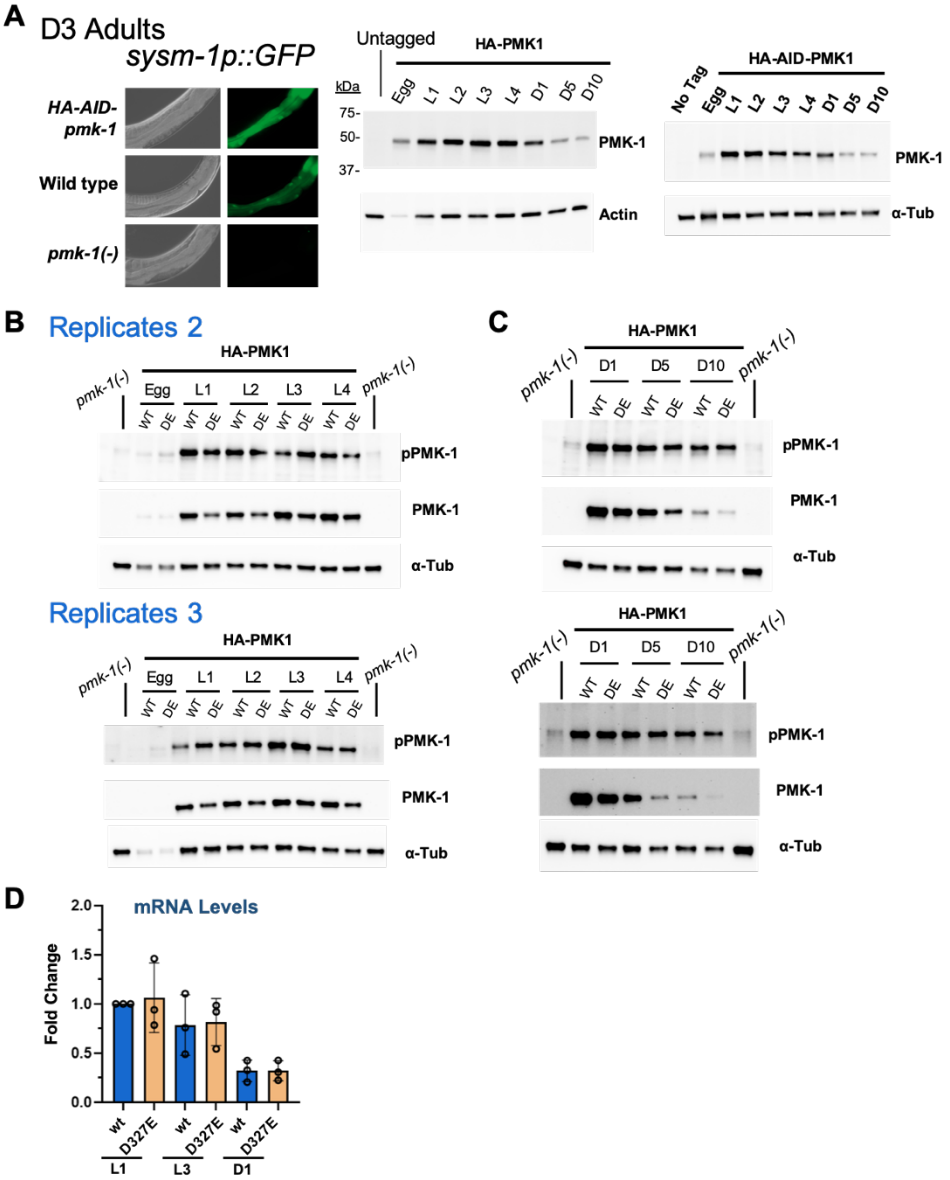


**Suppl. Fig. S2.** Replicates of PMK-1 expression with development and aging

**A)** PMK-1-dependent reporter gene induced with HA-AID-tagged PMK-1 at day3 adulthood.

**B-C)** Western blot analysis and quantitation of replicates for N-terminal tagged PMK-1 through development and aging. Loading controls, $\alpha$-tubulin or actin.

**D)** Quantitative RT-PCR of *pmk-1 (wt)* versus *pmk-1(D327E)* mRNAs normalized to *ama-1.* Standard deviation for 3 biological replicates (bars).

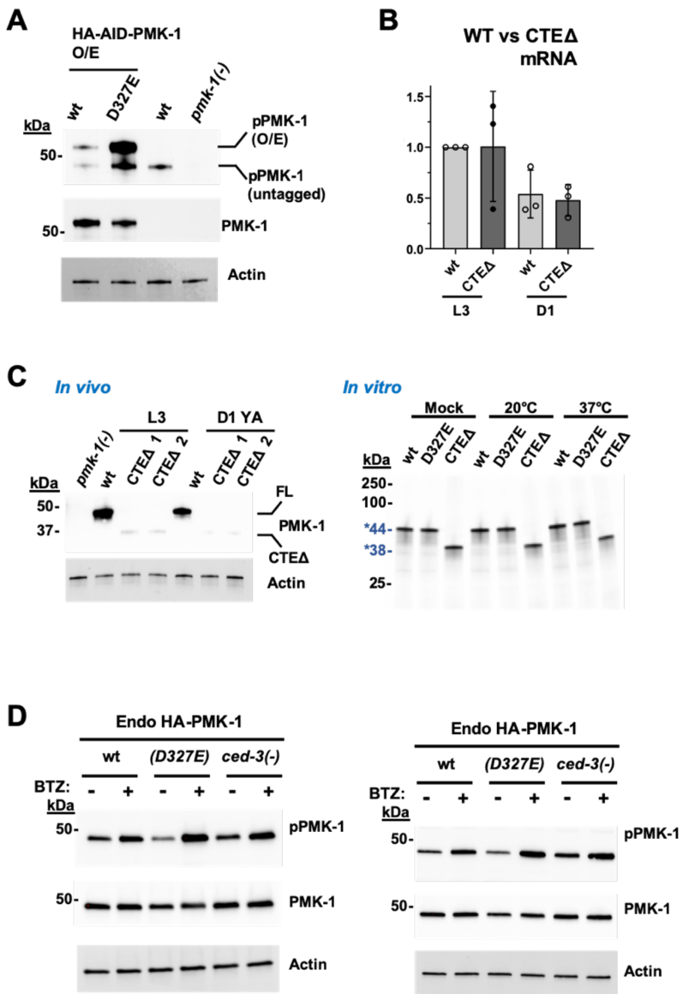


**Suppl. Fig. S3.** Overexpression replicate, mRNA levels, expression of CTE mutant, and Bortezomib replicates.

**A)** Replicate of overexpression.

**B)** Quantitative RT-PCR of *pmk-1 (wt)* versus *pmk-1(CTEΔ)* mRNAs normalized to *ama-1.* Standard deviation for 3 biological replicates (bars).

**C)** *In vivo* versus *in vitro* expression of CTE mutation.

**D)** Replicate of D327E overexpression enhancing pPMK-1 with Bortezomib treatment.

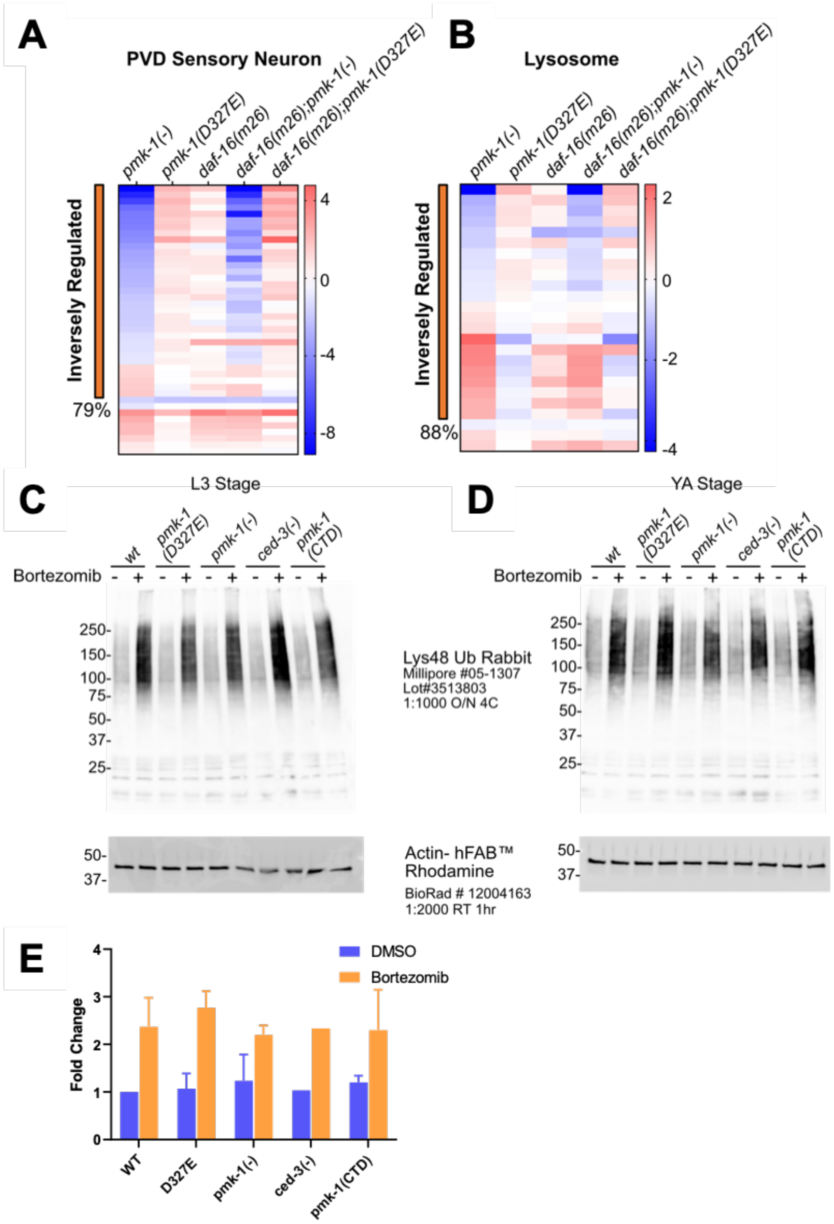


**Suppl. Fig. S4.** Heatmaps for *daf-16(m26)* allele and global K48-specific Ub linkage enrichments with Bortezomib treatment.

**A-B)** Sensory neuron and lysosome genes heatmap for *daf-16(m26)* allele.

**C-E)** Bortezomib treatment induced K48 linkages to comparable extent in *pmk-1(-)* null and *pmk-1(D327E)* cleavage resistant mutants.

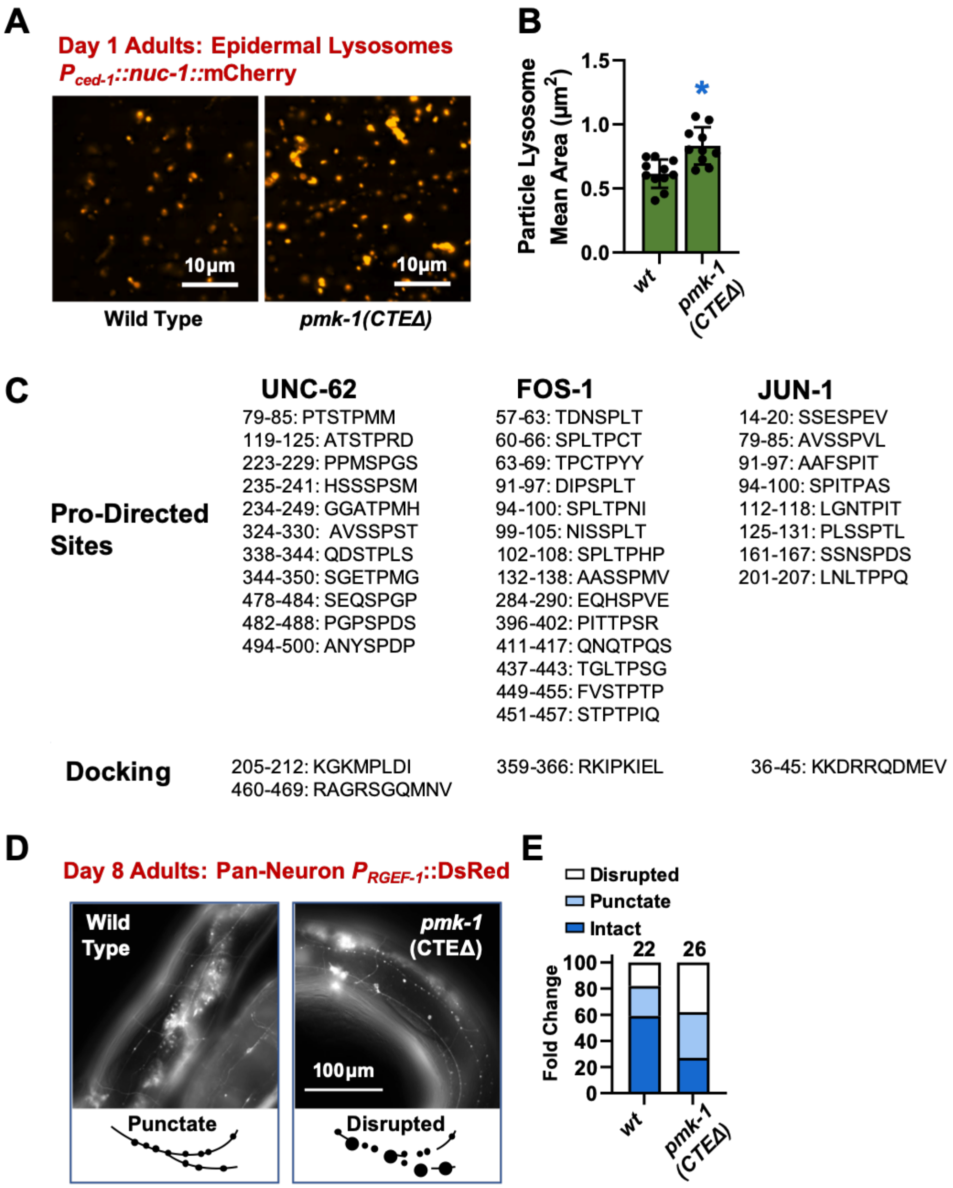


**Suppl. Figure S5.** Lysosome and neuron images of CTEΔ phenotypes

**A-B)** Day 1 adult lysosome morphology and quantitation in CTEΔ mutants compared to wild-type.

**C)** Motif analyses for putative phospho-sites and MAPK docking based on eukaryotic linear motif analysis (Eukaryotic Linear Motif database).

**D-E)** Day 8 images and quantitation of large mid-body neuronal morphologies in CTEΔ mutants compared to wild-type.

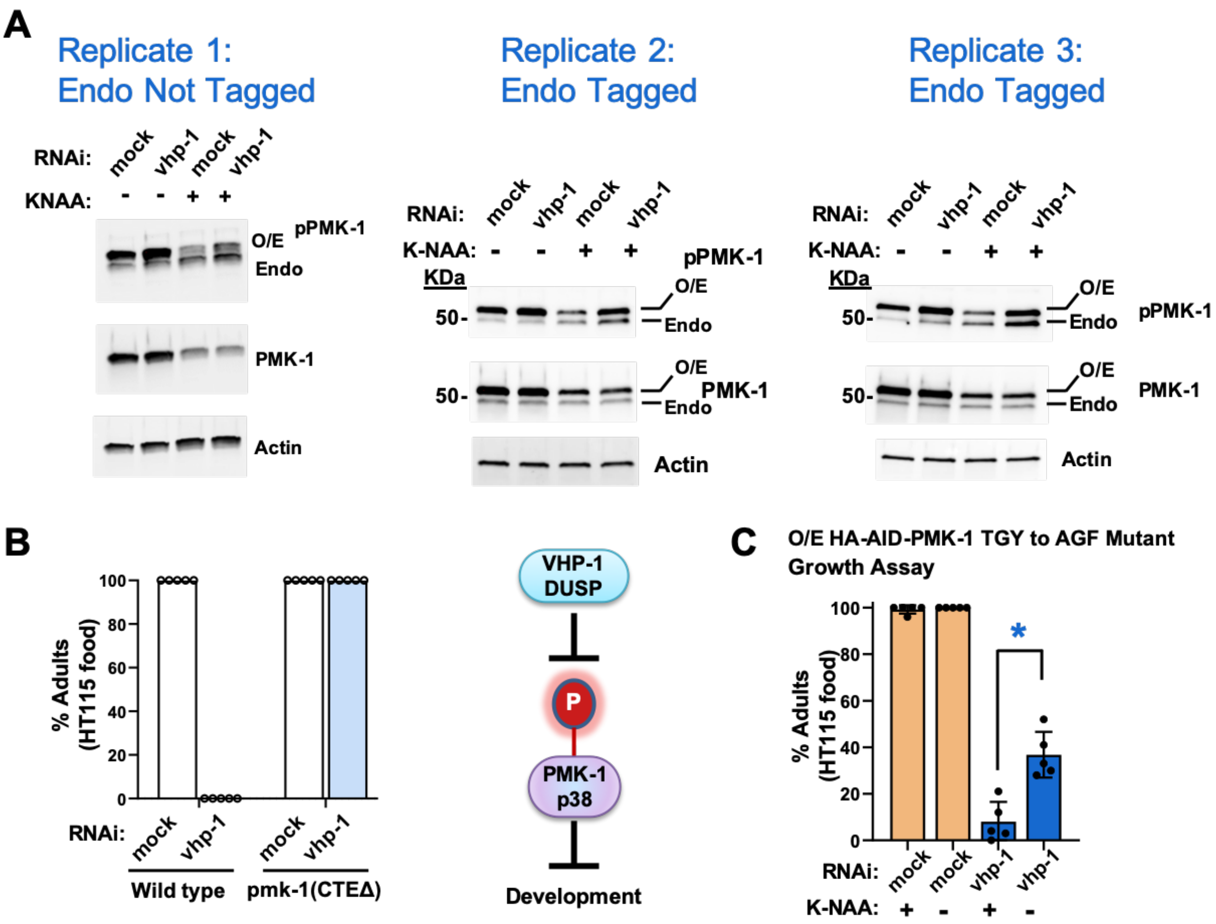


**Suppl. Fig. S6.** *pmk-1*-dependent growth stall, replicates for Western blot, and overexpression of *pmk-1* TGY to AGF mutant rescues growth on *vhp-1(RNAi)*

**A)** Western blot for conditions showing pPMK-1 and PMK-1 changes with *vhp-1(RNAi)* and auxin analog (K-NAA) treatments.

**B)** Developmental delay from *vhp-1* RNAi is *pmk-1*-dependent as shown by alleviation of the stall by loss of *pmk-1* function, consistent with previous findings by us and others.

**C)** Developmental rate assay for *pmk-1* TGY to AGF overexpression (O/E) mutant. Animals were treated with mock or *vhp-1(RNAi)* with (+) or without (-) auxin. Each dot corresponds to a plate with 25 to 50 animals, standard deviation (bars).

**Supplemental Tables**

**Supplemental Table S1.** *C. elegans* strains used in this study

**Supplemental Table S2.** sgRNA sequences for CRISPR mutagenesis used in this study

**Supplemental Table S3.** DNA repair templates used in this study

**Supplemental Table S4.** Statistics of aging assays

**Supplemental Table S5.** *pmk-1*-regulated genes

**Supplemental Table S6.** Transcription factor enrichment analysis

**Supplemental Table S7.** Mass-spectrometry for K48 polyUb IP
